# The most abundant cyst wall proteins of *Acanthamoeba castellanii* are three sets of lectins that bind cellulose and chitin and localize to distinct structures in cyst walls

**DOI:** 10.1101/496307

**Authors:** Pamela Magistrado-Coxen, Yousuf Aqeel, Angelo Lopez, John R. Haserick, Breeanna R. Urbanowicz, Catherine E. Costello, John Samuelson

## Abstract

*Acanthamoeba castellanii*, cause of keratitis and blindness, is an emerging pathogen because of its association with contact lens use. The cyst wall contributes to pathogenesis as cysts are resistant to sterilizing reagents in lens solutions and to antibiotics applied to the eye. We used transmission electron microscopy, as well as structured illumination microscopy and probes for cellulose and chitin, to show that purified cyst walls of *A. castellanii* retain an outer ectocyst layer, an inner endocyst layer, and conical ostioles that connect the layers. Mass spectrometry showed candidate cyst wall proteins are dominated by three families of lectins (named here Luke, Leo, and Jonah), each of which binds to microcrystalline cellulose and to a lesser degree chitin. A Jonah lectin, which has one choice-of-anchor A (CAA) domain, localizes to the ectocyst layer of mature cyst walls. Luke lectins, which have two or three carbohydrate-binding modules (CBM49), localize to the endocyst layer and ostioles. A Leo lectin, which has two domains with eight Cys residues each (8-Cys), also localizes to the endocyst layer and ostioles. In summary, the most abundant *A. castellanii* cyst wall proteins are three sets of lectins, which have carbohydrate-binding modules that are conserved (CBM49s of Luke), newly characterized (CAA of Jonah), or unique to *Acanthamoebae* (8-Cys of Leo). Despite their lack of common ancestry, Luke and Leo lectins both localize to the endocyst layer and ostioles, while the Jonah lectin localizes to the ectocyst layer.

**IMPORTANCE:** Fifty years ago, investigators identified cellulose in the *Acanthamoeba* cyst wall, which has an outer ectocyst layer, an inner endocyst layer, and conical ostioles that connect the layers. Here we show cyst walls also contain chitin and three large sets of cellulose- and chitin-binding lectins, which have distinct localizations. The *Acanthamoeba* cyst wall, therefore, is more complicated than cyst walls of *Entamoeba* and *Giardia* (causes of dysentery and diarrhea, respectively), which have a single layer, a single glycopolymer, and small sets of one or two lectins. In contrast, the *Acanthamoeba* cyst wall is far simpler than the walls of fungi and plants, which have multiple layers, numerous glycopolymers, and hundreds of proteins. In addition to providing a better understanding of the cell biology and biochemistry of the *Acanthamoeba* cyst wall, these studies may lead to diagnostic antibodies that bind to cyst wall proteins and/or therapeutics that target chitin.

## Introduction

*Acanthamoebae*, which include the genome project *A. castellanii* Neff strain, are soil protists named for acanthopods (spikes) on the surface of trophozoites (1). In immunocompetent persons, *Acanthamoebae* is a rare but important cause of corneal inflammation (keratitis), which is difficult to treat and so may lead to scarring and blindness (2-4). In immunosuppressed patients, *Acanthamoebae* may cause encephalitis (5). *Acanthamoeba* is an emerging pathogen, because 80 to 90% of infections are associated with contact lens use, which is growing worldwide (6, 7). Because water for washing hands may be scarce in places where the free-living protist is frequent, we recently showed that alcohols in concentrations present in hand sanitizers kill *A. castellanii* trophozoites and cysts (8, 9).

When *A. castellanii* trophozoites are deprived of nutrients in solution or on agar plates, they form cysts (10-12). Transmission electron microscopy (TEM) shows cyst walls have two microfibril-dense layers (outer ectocyst and inner endocyst), which are separated by a relatively microfibril-free layer (13). The endocyst and ectocyst layers are connected to each other by conical ostioles, through which the protist escapes during excystation (14).

The cyst wall of *A. castellanii* protects free-living protists from osmotic shock when exposed to fresh water, drying when exposed to air, or starvation when deprived of bacteria or other food sources. The cyst wall also acts as a barrier, sheltering parasites from killing by disinfectants used to clean surfaces, sterilizing agents in contact lens solutions, and/or antibiotics applied directly to the eye (15-17).

We are interested in the cyst wall proteins (CWPs) of *A. castellanii* for three reasons. First, although monoclonal antibodies to *A. castellanii* have been made, the majority react to trophozoites, and no CWPs have been molecularly identified (18-21). Indeed the only cyst-specific protein identified, which was named for its 21-kDa predicted size (CSP21), is unlikely to be a CWP, as it lacks a signal peptide (22). Second, *A. castellanii* and related species are the only human pathogens that contain cellulose in their wall (23-25). *Dictyostelium discoideum*, which also has cellulose in its walls, is not pathogenic (26). Third, the whole genome of *A. castellanii* predicts a chitin synthase, a chitinase, and two chitin deacetylases, suggesting the possibility that chitin and chitin-binding proteins are also present in the cyst wall (1, 27-29).

Our experimental design was simple. We used transmission electron microscopy (TEM), as well as structured illumination microscopy (SIM) and probes for glycopolymers, to judge the intactness and cleanliness of purified *A. castellanii* cyst walls (30). We used mass spectrometry to identify candidate CWPs, which were compared to proteins present in walls of other protists, bacteria, fungi, and plants (31). We used SIM to localize representative CWPs, each of which was tagged with a green fluorescent protein (GFP) and expressed under its own promoter in cyst walls of stably transfected protists (32, 33). We also determined whether each CWP, which was made as a maltose-binding protein (MBP) in the periplasm of bacteria, binds to microcrystalline cellulose and/or chitin beads (34).

## RESULTS

### TEM and SIM showed purified *A. castellanii* cyst walls contained distinct endocyst and ectocyst layers, as well as ostioles

Cyst wall preparations were made by sonicating cysts and separating walls from cellular contents by density centrifugation and retention on a membrane with 8 µm-diameter pores. For TEM, intact cysts and purified cyst walls were frozen under high pressure, and fixatives were infiltrated at low temperature (35). Purified cyst walls had intact ectocyst and endocyst layers, as well as conical ostioles that link them (Fig. 1) (13). The purified walls were missing amorphous material that fills the space between the inner aspect of the cyst wall and the plasma membrane of the trophozoite inside.

**FIG 1.**
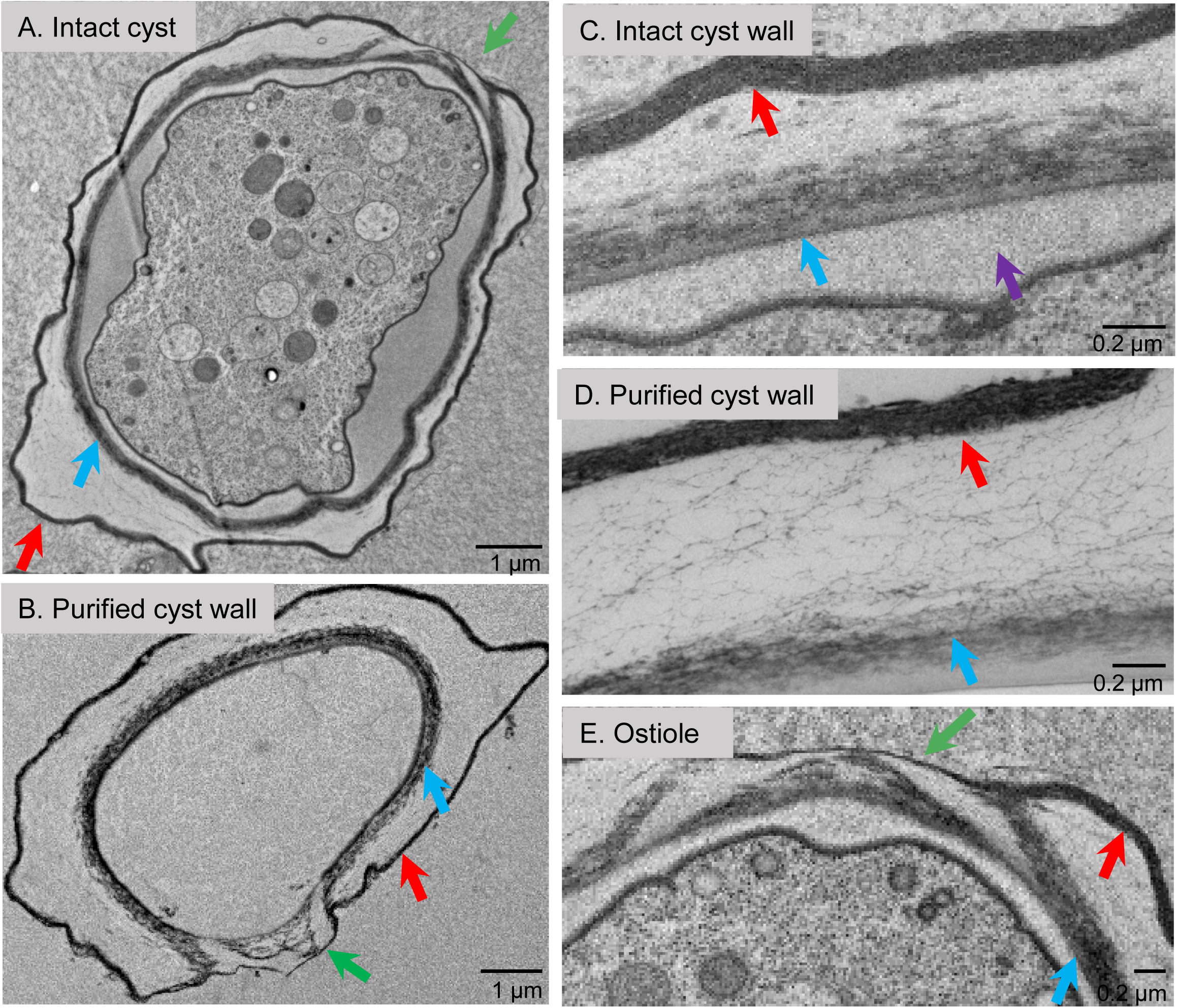
TEM showed purified *A. castellanii* cyst walls retained endocyst and ectocyst layers and ostioles. A, B. Intact cysts and purified cyst walls had an outer ectocyst layer (red arrows), an inner endocyst layer (blue arrows), and ostioles (green arrows) that connect the layers. Endocyst and ectocyst layers had the same appearance in intact cysts (C) and purified cyst walls (D). Purified cyst walls were missing amorphous material (purple arrow) between the wall and the plasma membrane of intact cysts. E. At the edge of the ostiole of an intact cyst, the endocyst layer bifurcated, and the outer branch met the ectocyst layer. In the center of the ostiole, the ectocyst layer formed a narrow cap over the inner branch of the endocyst layer. Scale bars as marked on micrographs.

For SIM, we used probes that bind chitin (wheat germ agglutinin, WGA) and β-1,3 and β-1,4 polysaccharides (calcofluor white, CFW) in the walls of fungi and cysts of *Entamoeba* (36-39). CFW, a fluorescent brightener, has also been used to diagnose *Acanthamoeba* cysts in eye infections (40). In addition, we made a glutathione-S-transferase (GST) fusion-protein, which contains the N-terminal CBM49 of a candidate CWP of *A. castellanii* (Fig. S1 and Excel file S1) (41). The GST-AcCBM49 expression construct was designed to replicate that used to determine the carbohydrate binding properties of SlCBM49, which is a C-terminal carbohydrate-binding module of the *Solanum lycopersicum* (tomato) cellulase SlGH9C (42). In both intact cysts and purified cyst walls, GST-AcCBM49 predominantly labeled the ectocyst layer, WGA highlighted the ostioles, and CFW stained the endocyst layer (Fig. 2). In summary, TEM and SIM both showed that ectocyst and endocyst layers, as well as ostioles, were intact in purified cyst walls, which were relatively free of cellular material.

**FIG 2.**
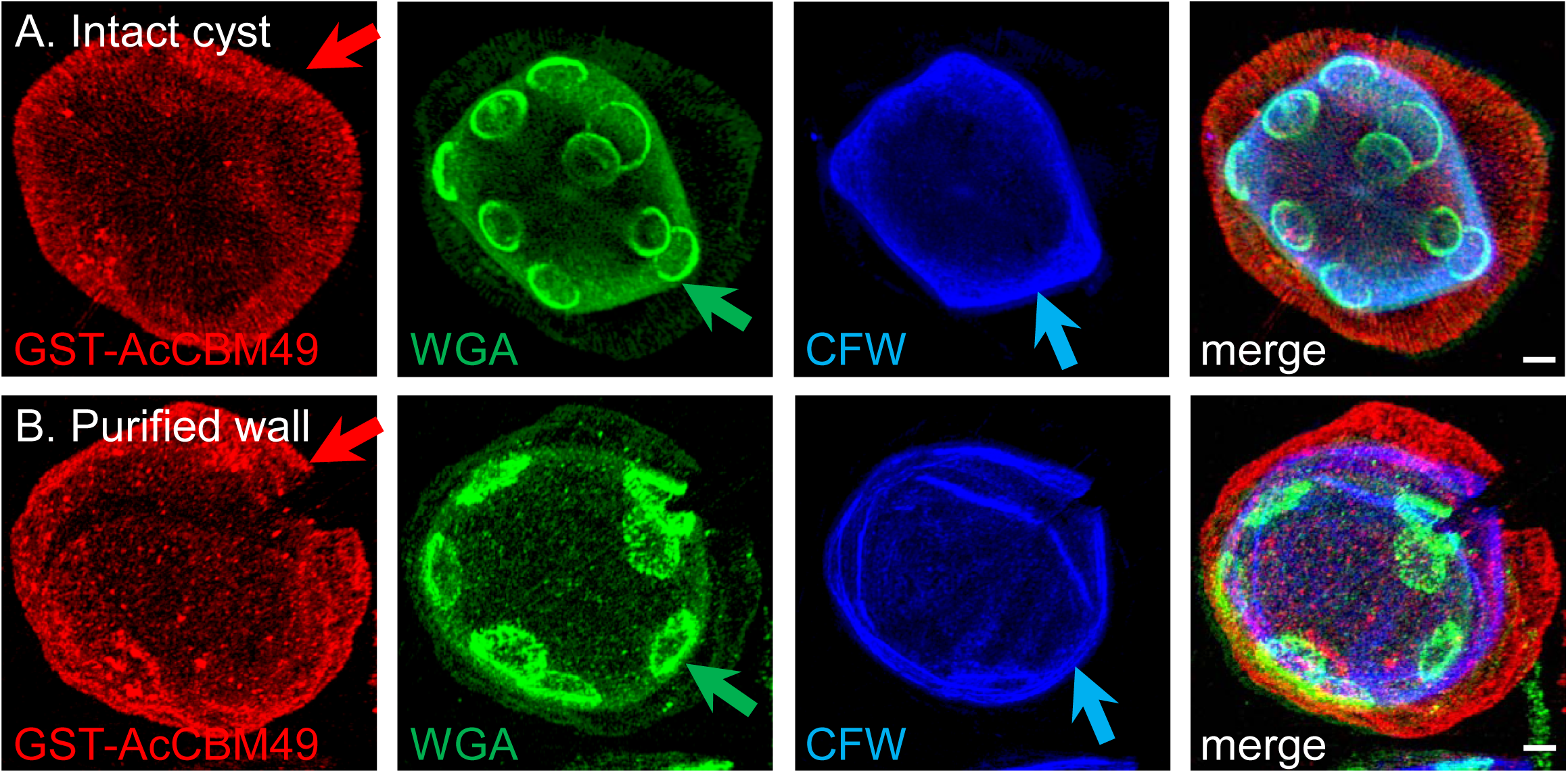
SIM also showed purified *A. castellanii* cyst walls retained distinct ectocyst layer (red arrow) and endocyst layer (blue arrow), as well as ostioles (green arrow). The ectocyst layer of intact cysts (A) and purified cyst walls (B) labeled red with GST-AcCBM49; the edges of ostioles labeled green with WGA; and the endocyst layer labeled blue with CFW. Scale bars are 2 µm.

### Mass spectrometry showed candidate CWPs of *A. castellanii* were encoded by multigene families and contain tandem repeats of short domains (CBM49, 8-Cys, and CAA)

Trypsin treatment of purified *A. castellanii* cyst walls, which was followed by liquid chromatography mass spectrometry (LC-MS/MS) analysis of the released peptides, gave similar results in five biological experiments (Table 1 and Excel file S1). While some proteins remained in cyst walls after trypsin digestion, their identities were similar to those detected in the soluble fractions by in gel-digests with trypsin or chymotrypsin. Candidate CWPs with the most unique peptides identified by LC-MS/MS belonged to three families, which we named Luke, Leo, and Jonah lectins, because each bound to cellulose +/- chitin (see below). Although it was impossible to draw a line that separates actual CWPs from contaminating proteins, secreted proteins with 18+ unique peptides included six Leo lectins, four Luke lectins, and three Jonah lectins. The vast majority of proteins with <18 unique peptides were predicted to be cytosolic (including CSP21) and so were likely intracellular contaminants of cyst wall preparations. The exception to this hypothesis, we think, were additional Luke, Leo, and Jonah lectins, which are most likely less abundant CWPs. For readers interested in cytosolic proteins of *A. castellanii*, we have added Excel file S2, which contains all the mass spectrometry data including a “dirty” cyst wall preparation that was generated without using the Percoll gradient or porous filter.

**Table 1.** Candidate CWPs identified by mass spectrometry.

|  | ID | # Unique peptides | Coverage (%) | Mass (kDa) |
| --- | --- | --- | --- | --- |
| **Jonah lectins** |  |  |  |  |
| three CAAs | ACA1_157320 | 147 | 38 | 146 |
| one CAA | ACA1_164810 | 83 | 56 | 58 |
|  | ACA1_261530 | 18 | 23 | 55 |
|  | ACA1_133400 | 9 | 24 | 44 |
|  | ACA1_377440 | 6 | 11 | 47 |
| **Luke lectins** |  |  |  |  |
| three CBM49s | ACA1_245650 | 72 | 74 | 44 |
|  | ACA1_160160 | 8 | 25 | 43 |
|  | ACA1_187760 | 7 | 25 | 42 |
|  | ACA1_252830 | 6 | 19 | 44 |
|  | ACA1_031530 | 6 | 21 | 43 |
|  | ACA1_253650 | 5 | 20 | 42 |
|  | ACA1_253500 | 5 | 19 | 42 |
|  | ACA1_061050 | 3 | 12 | 43 |
|  | ACA1_287530 | 2 | 14 | 43 |
| two CBM49s | ACA1_377670 | 78 | 68 | 29 |
|  | ACA1_096300 | 47 | 77 | 28 |
|  | ACA1_246110 | 22 | 70 | 27 |
| **Leo lectins** |  |  |  |  |
| two 8-Cys domains | ACA1_074730 | 34 | 82 | 20 |
|  | ACA1_351320 | 24 | 44 | 20 |
|  | ACA1_394030 | 24 | 44 | 20 |
|  | ACA1_394280 | 24 | 36 | 24 |
|  | ACA1_083920 | 19 | 68 | 20 |
|  | ACA1_394560 | 1 | 10 | 19 |
| two 8-Cys + TKH | ACA1_188350 | 21 | 20 | 59 |
|  | ACA1_374130 | 7 | 20 | 52 |
|  | ACA1_188550 | 7 | 15 | 46 |
|  | ACA1_188370 | 6 | 9 | 68 |
|  | ACA1_116240 | 5 | 18 | 56 |
|  | ACA1_365840 | 3 | 18 | 44 |
|  | ACA1_117050 | 3 | 33 | 36 |
|  | ACA1_096640 | 2 | 27 | 37 |

Luke lectins were comprised of an N-terminal signal peptide, followed by two or three CBM49s that were separated by Ser- and Pro-rich spacers (Figs. 3 and S2) (27, 42-45). The N-terminal CBM49 of Luke lectins contained three conserved Trp resides conserved in SlCBM49 from tomato. A fourth conserved Trp is present in the CBM49 of *D. discoideum* cellulose-binding proteins (46). The other CBM49s (middle and/or C-terminal) of Luke lectins had two conserved Trp residues. Luke lectins were acidic (pI 5 to 6) and had formula weights (FWs) from 27 to 29-kDa (two CBM49s) or 42 to 44-kDa (three CBM49s). There were no predicted transmembrane helices or glycosylphosphatidylinositol anchors in the Luke or Leo lectins (47, 48). LC-MS/MS of the released cell wall peptides identified at least one unique peptide corresponding to all 12 genes encoding Luke lectins, although the number of unique peptides varied from 78 to two (Table 1 and Excel file S1). In general, Luke lectins with two CBM49s had more unique peptides than Luke lectins with three CBM49s. One to four unique peptides were derived from three CBM49-metalloprotease fusion-proteins, which consisted of an N-terminal signal peptide followed by a single CBM49 with four conserved Trp residues and a metalloprotease (ADAM/reprolysin subtype) with a conserved catalytic domain (HEIGHNLGGNH) (43). We used a Luke(2) lectin (ACA1_377670) with two CBM49s to perform RT-PCR, make rabbit anti-peptide antibodies, and make maltose-binding protein (MBP)- and green fluorescent protein (GFP)-fusions (Figs. 3 and S1) (33, 34). We also used a Luke(3) lectin (ACA1_245650) with three CBM49s to make a GFP-fusion (Fig. S2).

**FIG 3.**
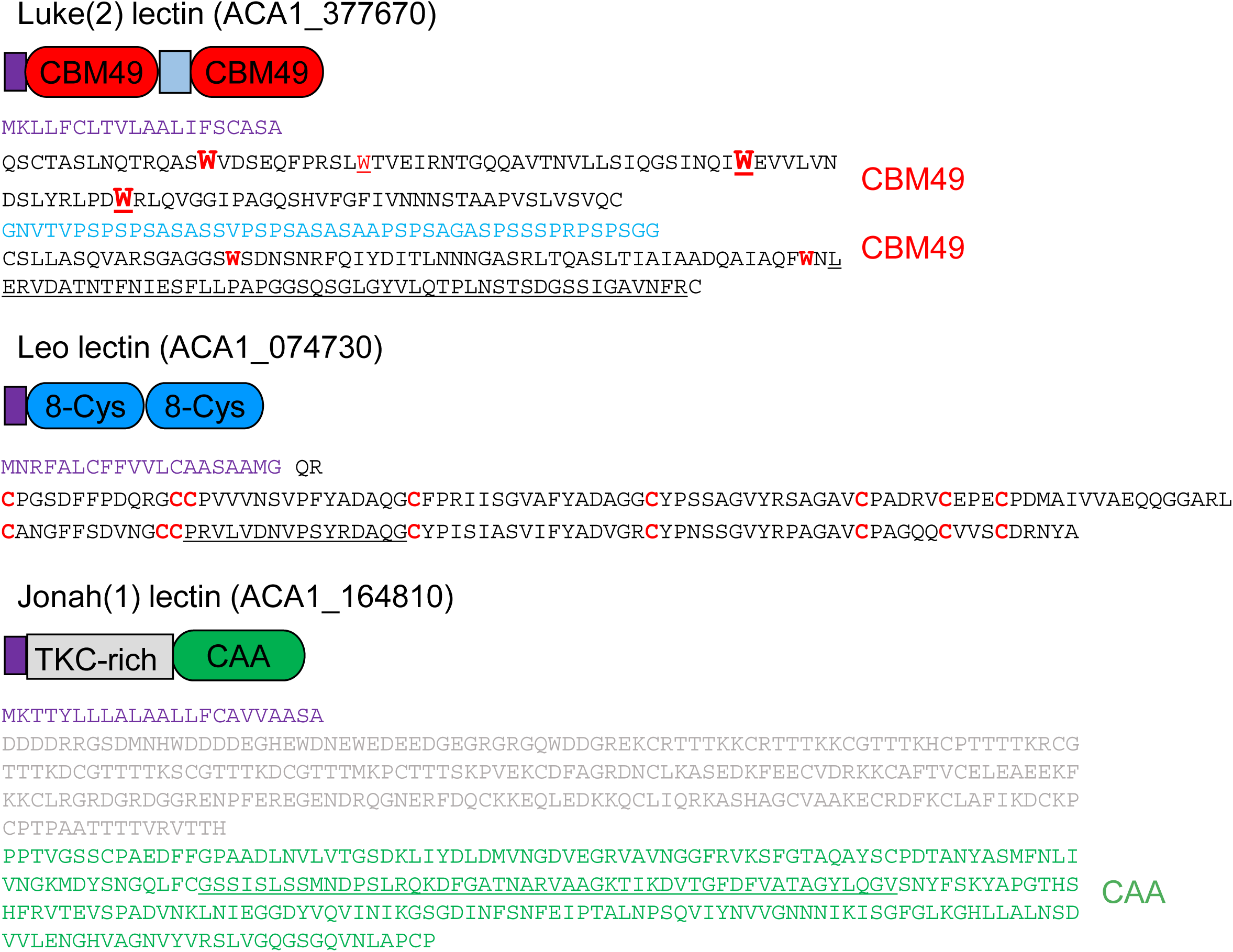
Representative proteins of abundant families of CWPs contained two CBM49s (Luke(2) lectin), two 8-Cys domains (Leo lectin), or one CAA domain (Jonah(1) lectin). The Luke(2) lectin had an N-terminal signal peptide (purple) and two CBM49s separated by short Ser- and Pro-rich spacers (light blue). The N-terminal CBM49 contained four Trp residues (red Ws), three of which were conserved in a C-terminal CBM49 of a tomato cellulase (larger font) and three of which were conserved in a single CBM49 of *Dictyostelium* spore coat proteins (underlined). In contrast, the C-terminal CBM49 had two conserved Trp residues. The Leo lectin had a signal peptide and two unique domains (dark blue) containing eight Cys residues each (red Cs). The Jonah(1) lectin had a signal peptide, a Thr-, Lys-, and Cys-rich domain (tan), and a single CAA domain (green). Peptides used to immunize rabbits are underlined. Representative Luke(3) lectin with three CBM49s, Leo(TRH) lectin with a long Thr-, Lys-, and His-rich spacer, and Jonah(3) lectin with three CAA domains are shown in Fig. S2.

Leo lectins had an N-terminal signal peptide followed by two repeats of a unique 8-Cys domain, some of which were separated by a long Thr-, Lys-, and His-rich spacer (Figs. 3 and S2). Leo lectins without a spacer were acidic (pI ∼4.8) and had FWs from 19 to 24-kDa, while Leo lectins with the TKH-rich spacer were basic (pI ∼8.3) and had FWs from 36- to 59-kDa. Leo lectins were encoded by 16 genes, of which 14 proteins were identified by our LC-MS/MS analysis. While the number of unique peptides varied from 34 to one, Leo lectins without the spacer generally had more unique peptides than Leo lectins with the TKH-rich spacer. We used a Leo lectin with no spacer (ACA1_074730) to perform RT-PCR, make rabbit anti-peptide antibodies, and make MBP- and GFP-fusions (Fig. S1).

Jonah lectins had an N-terminal signal peptide followed by one or three choice-of-anchor A (CAA) domains (Figs 3. and S2) (43). The binding activity of the CAA domain, which is adjacent to a collagen-binding domain in a microbial surface component recognizing the adhesive matrix molecule (MSRAMM) of *Bacillus anthracis*, was not known (49). Jonah lectins with a single CAA domain were acidic (pI ∼6), had a FW from 44 to 58-kDa and had an N-terminal Thr-, Lys-, and Cys-rich domain. Jonah lectins with three CAA domains were basic (pI ∼8.8), had a FW of ∼146-kDa, and contained Ser- and Pro-rich spacers between CAA domains, as well as and hydrophobic regions that may be transmembrane helices (48). Jonah lectins were encoded by eight genes, of which five were identified by our LC-MS/MS analysis based on one to 147 unique peptides. We used a Jonah lectin (ACA1_164810) with a single CAA domain to perform RT-PCR, make rabbit anti-peptide antibodies, and make MBP- and GFP-fusions (Fig. S1).

Other secreted proteins with 18+ unique peptides detected by LC-MS/MS, which are candidate CWPs, included a laccase with three copper oxidase domains (ACA1_068450), a protein with a C-terminal ferritin-like domain (ACA1_292810), a Kazal-type serine protease inhibitor (ACA1_291590), a conserved uncharacterized protein (ACA1_068630), and a protein unique to *A. castellanii* (ACA1_145900) (43, 44, 50). Interestingly, a bacterial laccase has been shown to bind cellulose (51). There were also three serine proteases, which have been localized to the secretory system of encysting organisms (52).

These results suggested that the most abundant candidate CWPs of *A. castellanii* contain tandem repeats of conserved domains (CBM49 in Luke lectins and CAA in Jonah lectins) or a unique domain (8-Cys in Leo lectins). Peptides corresponding to nearly all members of each gene family were detected by mass spectrometry, although the relative abundances of unique peptides for each CWP varied by more than an order of magnitude, suggesting marked differences in gene expression. Because it was not possible to separate cyst walls into component parts (endocyst and ectocyst layers and ostioles) prior to LC-MS/MS analysis of tryptic peptides, we used SIM and GFP-tags to localize representative members of each family of CWPs in cyst walls of transfected *A. castellanii* (see below).

### Origins and diversity of genes that encode Luke, Leo, and Jonah lectins

Leo lectins, which had two domains with 8-Cys each, appeared to be unique to *A. castellanii*, as no homologs were identified when BLAST analysis were performed using the nonredundant (NR) database at NCBI (https://www.ncbi.nlm.nih.gov/) (28). The origin of genes encoding Luke lectins was difficult to infer, because its CBM49s showed only a 31% identity over a short (77-amino acid) overlap with a predicted cellulose-binding protein of *D. discoideum* (expect value of BLASTP was just 7e-05) (28, 46). In contrast, the CAA domain of Jonah lectins appeared to derive from bacteria by horizontal gene transfer (HGT), as no other eukaryote contained CAA domains, and there was a 28% identity over a bigger (263-aa) overlap with a choice-of anchor A family protein of *Saccharibacillus sp. O16* (5e-12) (1, 28). The *A. castellanii* laccase (also known as copper oxidase), whose signals were abundant in the mass spectra, was likely the product of HGT from bacteria, as there was a 44% identity over a large (526-aa) overlap with a copper oxidase of *Caldicobacteri oshimai* (6e-135) (50). The uncertainty was based upon the presence of similar enzymes in plants, one of which (*Ziziphus jujube*) showed a 39% identity over a 484-aa overlap (4e-101) with the *A. castellanii* laccase.

No pairs of genes within each lectin family were syntenic, suggesting that individual genes rather than clusters of genes were duplicated. With the exception of two Luke lectins (ACA1_253500 and ACA1_253650) that were 98% identical and two Leo lectins (ACA1_074770 and ACA1_083920) that were 85% identical, members of each family of CWPs differed in amino acid sequence by >40%. Genes that encode CWPs also varied in the number of introns (zero to two in Luke, two to four in Leo, and zero to 24 in Jonah). Searches of genomic sequences of 11 strains of *Acanthamoebae*, deposited in AmoebaDB without protein predictions by Andrew Jackson of the University of Liverpool, using TBLASTN and sequences of representative Luke, Leo, and Jonah lectins localized in the next section, showed four results (28, 29). First, although stop codons were difficult to identify using this method, all 11 strains appeared to encode each CWP. Second, most strains showed 100 to 200-amino acid stretches of each CWP that were 80 to 90% identical to the *A. castellanii* Neff strain studied here. These stretches did not include low complexity spacers, which are difficult to align. Third, some of the strains showed greater differences from the Neff strain in each CWP, consistent with previous descriptions of *Acanthamoeba* strain diversity based upon 18S rDNA sequences (53). Fourth, while coding sequences and 5’ UTRs were well conserved, introns were very poorly conserved, with the exception of branch-point sequences.

In summary, genes encoding Jonah lectin and laccase likely derived by HGT, while genes encoding Leo lectins appeared to originate within *Acanthamoeba*. Although CBM49s of Luke lectins shared common ancestry with plants and other Amoebazoa, their precise origin was not clear. For the most part, gene duplications that expanded each family within the *Acanthamoeba* genome occurred a long time ago, as shown by big differences in amino acid sequences of paralogous proteins and variations in the number of introns. Still, the set of Luke, Leo, and Jonah lectins identified by mass spectrometry and the sequence of representative CWPs localized in the next section both appeared to be conserved among 11 sequenced isolates of *Acanthamoebae*.

### SIM showed a representative Jonah lectin was present in the ectocyst layer of mature cyst walls, while representative Luke and Leo lectins were present in the endocyst layer and ostioles

To localize candidate CWPs, we expressed a Leo lectin with no spacer and a Jonah(1) lectin with a single CAA domain, each with a GFP-tag under its own promoter (446- and 571-bp of the 5’ UTR, respectively) in transfected trophozoites of *A. castellanii*, using an episomal vector that was selected with G418 (Fig. S1) (32, 33). We also expressed a Luke(2) lectin with two CBM49s and a Luke(3) lectin with three CBM49s, each with a GFP-tag under its own promoter (486- and 500-bp of the 5’ UTR, respectively). GFP-tagged candidate CWPs expressed under their own promoter were absent in the vast majority of log-phase trophozoites, which divided but did not become cysts (data not shown). GFP-tagged CWPs were present in small numbers in trophozoites in stationary cultures, where a few organisms have begun to encyst spontaneously. In contrast, the vast majority of mature cysts contained GFP-tagged CWPs in their walls.

Jonah(1)-GFP was present in the ectocyst layer of mature cyst walls, which were labeled with WGA and CFW (Fig. 4A). In contrast, Leo-GFP, Luke(2)-GFP, and Luke(3)-GFP were each present in the endocyst layer and sharply outlined the ostiole (Figs. 4B to 4D). These results showed that ∼500 bp of the 5’ UTR was sufficient to cause encystation-specific expression of GFP-tagged representatives of Luke, Leo, and Jonah lectins. While the Jonah lectin localized to the ectocyst layer, Luke and Leo lectins, which do not share common ancestry, both localized to the endocyst layer and ostioles. Finally, Luke lectins with two or three CBM49s localized to the same place.

**FIG 4.**
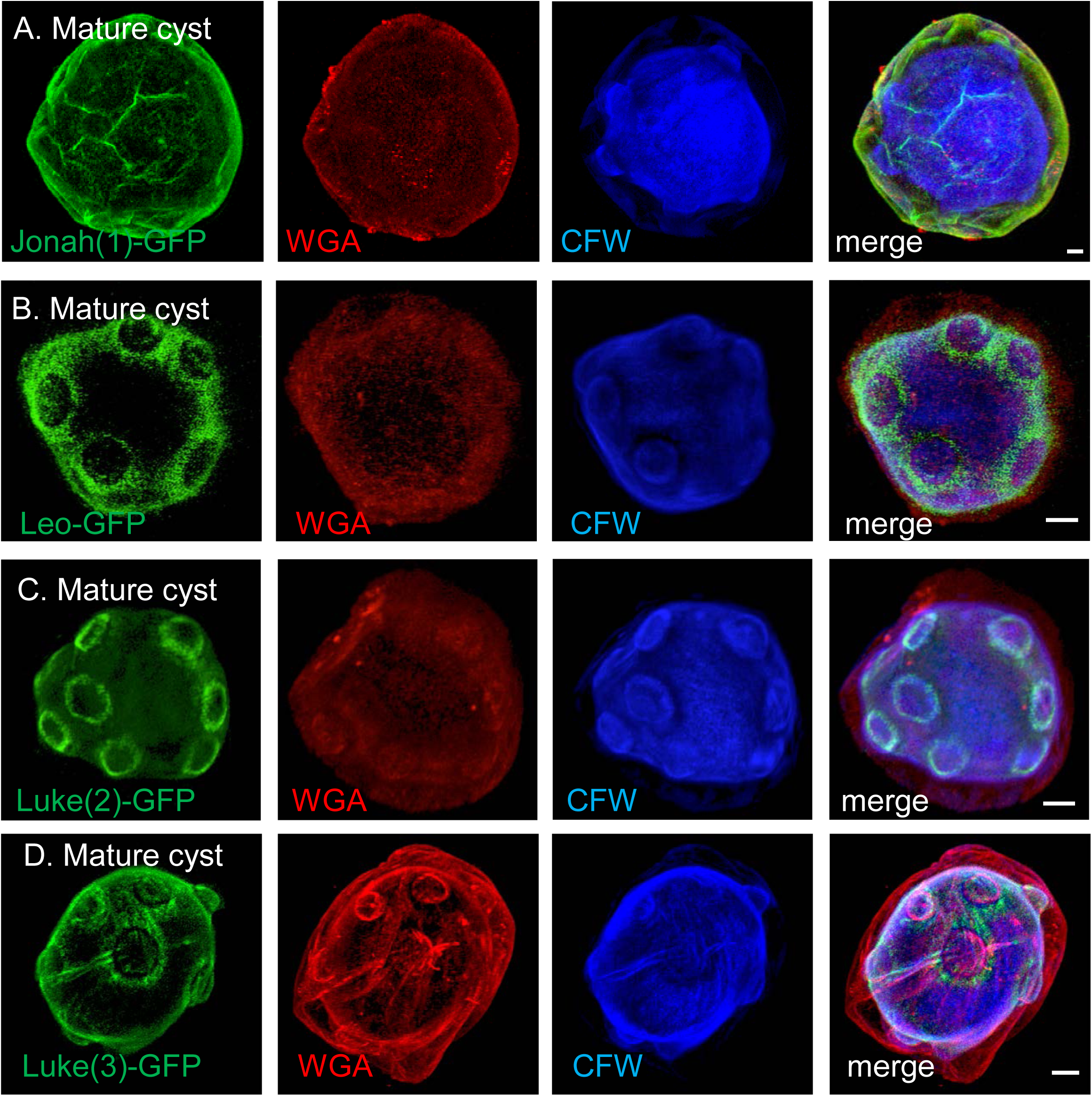
SIM showed a representative Jonah lectin localized to the ectocyst layer of mature cyst walls, while representative Luke lectins and Leo lectins localized to the endocyst layer and ostioles. A. A Jonah(1) lectin with a single CAA domain, which was tagged with GFP and expressed under its own promoter in transfected *A. castellanii*, localized to the ectocyst layer of the wall of mature cysts. WGA labeled both ectocyst and endocyst layers, while CFW labeled the endocyst layer and ostioles. B. Leo-GFP with two 8-Cys domains was also present in the endocyst layer and ostioles. C. Luke(2)-GFP with two CBM49s was present in the endocyst layer and sharply outlined conical ostioles of mature cysts. D. Luke(3) with three CBM49s was also present in the endocyst layer and ostioles. A to D. Scale bars are 2 µm.

We think the timing of expression and locations of the GFP-tagged CWPs in cyst walls were accurate for the following reasons. First, RT-PCR showed that mRNAs of representative Luke, Leo, and Jonah lectins, as well as cellulose synthase (ACA1_349650), were absent or nearly absent from trophozoites but were present during the first three days of encystation (Fig. S3). In contrast, glyceraldehyde 3-phosphate dehydrogenase (GAPDH), which catalyzes the sixth step in glycolysis, was expressed by both trophozoites and encysting *A. castellanii* (33). Second, monospecific, polyclonal rabbit antibodies to a 50-amino acid peptide of a representative Jonah lectin and a 16-amino acid peptide of a representative Leo lectin bound to Western blots of proteins from cysts but not from trophozoites (Fig. S4). We were unable to generate rabbit antibodies to the Luke lectin, using methods that worked to make antibodies to Jonah and Leo lectins. None of the rabbit anti-peptide antibodies was useful for localizing CWPs by SIM. Third, GFP-tagged CSP21 expressed under its own promoter was present in cytosolic accumulations of mature cysts (Fig. S5) (22, 54). As CSP21 is homologous to universal stress proteins and lacks an N-terminal signal peptide, its presence in the cytosol after nutrient deprivation was expected (43, 55). Fourth, a GFP-fusion protein (SP-GFP), which was appended with an N-terminal signal peptide from Luke lectin and expressed under a GAPDH promoter, localized to secretory vesicles of cysts but not to cyst walls (Fig. S5) (33).

### Luke, Leo, and Jonah lectins all bound to microcrystalline cellulose, while binding of CWPs to chitin beads was variable

To test the binding of representative CWPs to commercially available glycopolymers, we made MBP-CWP fusion-proteins in the periplasm (secretory compartment) of *E. coli* (34). The targets were microcrystalline cellulose (used to characterize binding activities of GST-SlCBM49 from tomato cellulase) and chitin beads (used to characterize myc-tagged Jacob and Jessie lectins of *Entamoeba histolytica*) (42, 56). Western blots anti-MBP antibodies showed MBP-Luke(2), which was partially degraded, bound to microcrystalline cellulose and somewhat less well to chitin beads (Fig. 5). MBP-Jonah(1) also bound to microcrystalline cellulose and chitin beads. MBP-Leo bound less completely to microcrystalline cellulose and weakly at best to chitin beads. MBP alone (negative control) did not bind to microcrystalline cellulose or chitin beads. These results suggested each CWP is binding to glycopolymers in the cyst wall, but they did not explain why Jonah localized to the ectocyst layer, while Luke and Leo localized to the endocyst layer and ostioles.

**FIG 5.**
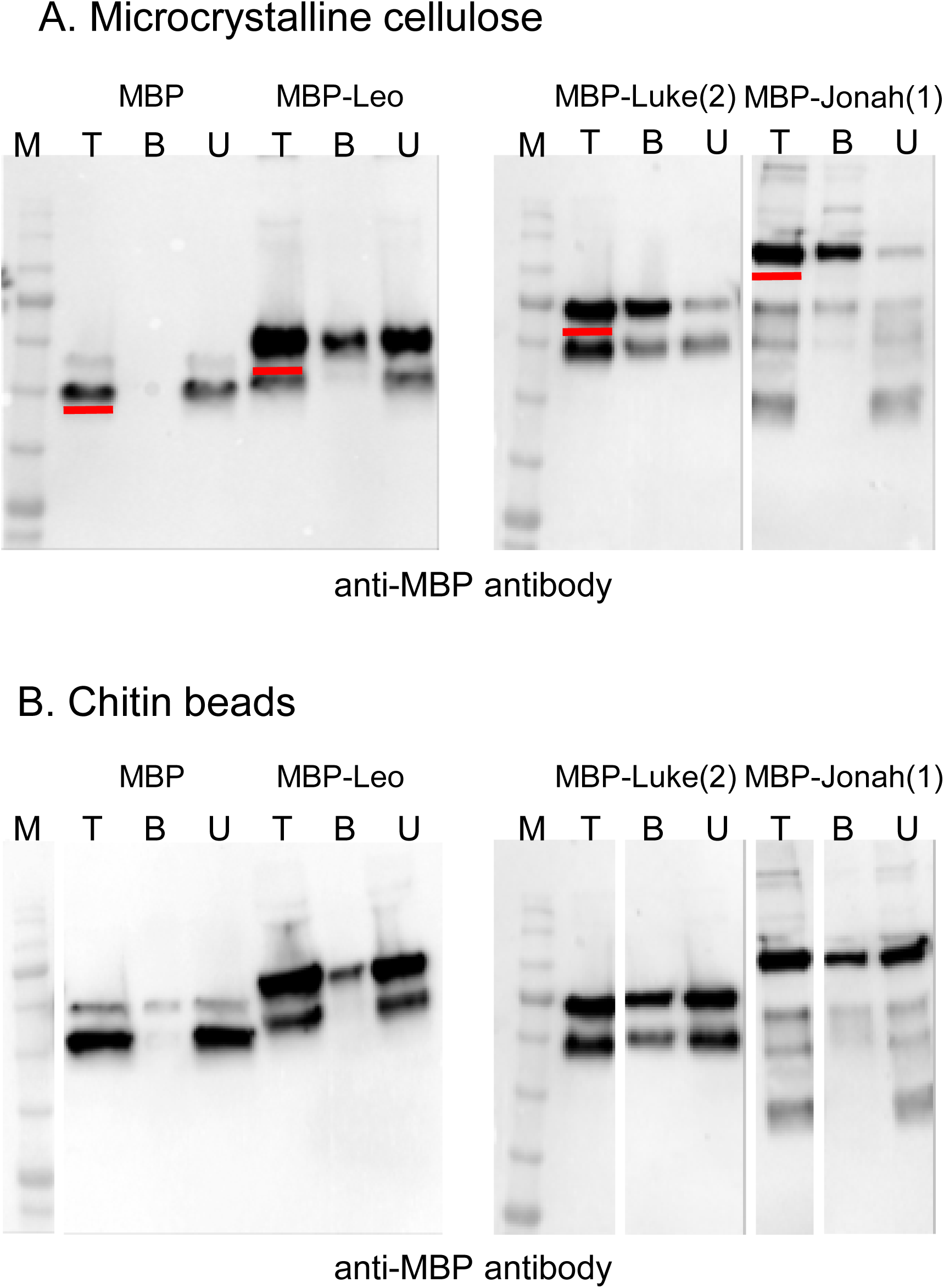
Western blots showed MBP-fusions to Luke, Leo, and Jonah lectins all bound to microcrystalline cellulose, while binding to chitin beads was variable. MBP-CWP fusions and MBP alone, made as recombinant proteins in the periplasm of bacteria, were incubated with microcrystalline cellulose (A) or chitin beads (B). Total proteins (T), bound proteins (B), and unbound proteins (U), as well as molecular weight markers (M), were run on SDS-PAGE, transferred to PVDF membranes, and detected with an anti-MBP reagent. Full-length products in total fractions are underlined in red. MBP-Leo partially bound to microcrystalline cellulose and bound weakly, if at all to chitin. MBP-Luke(2) and MBP-Jonah(1) each bound more completely to cellulose than to chitin, while MBP alone (negative control) did not bind to cellulose or chitin.

## Discussion

### Advantages of studying *Acanthamoeba* cyst walls

*A. castellanii* has a number of properties that make it possible to quickly identify and begin to characterize its CWPs. The protist grows axenically in medium without serum or vitamins and synchronously encysts when placed on non-nutrient agar plates (10, 11). Cyst walls separate cleanly from cellular contents without losing essential structures (ectocyst and endocyst layers and ostioles), which we show are readily visualized with SIM using probes for wall glycopolymers (GST-AcCBM49, WGA, and CFW) (13, 30, 36-39). The most abundant CWPs belong to three protein families that contain tandem repeats of CBM49s (Luke lectins), 8-Cys domains (Leo lectins), or CAA domains (Jonah lectins). Stable expression of GFP-tagged proteins under their own promoters allows localization of candidate CWPs during cyst wall development using SIM (32, 33). Efficient expression of recombinant proteins in the periplasm (MBP-fusions) of bacteria makes it possible to assay the ability of CWPs to bind microcrystalline cellulose or chitin beads, by using techniques to characterize other protist cyst wall lectins and carbohydrate-binding modules (34, 42, 56-58).

The ease of studying cyst walls of *A. castellanii* is in contrast to studies of cyst walls of *E. histolytica*, which do not encyst *in vitro* and so are modeled by cysts of the reptilian pathogen *E. invadens* (38, 56). While *Giardia* cysts made in mice excyst well, cysts made *in vitro* do not (57). *Toxoplasma* forms walled oocysts in cats, while *Cryptosporidium* forms oocysts in cows, mice, or gnotobiotic pigs (58-60). Oomycetes (water molds that cause potato blight) also have cellulose and chitin in their walls, but the amount of chitin is very small, and β- 1,3-glucan is also present (61). The spore coat of *Dictyostelium discoideum*, a cousin of *A. castellanii*, contains cellulose and a heteropolysaccharide of galactose and GalNAc, but it lacks chitin (26, 62).

Properties of *A. castellanii* that make it challenging to study its cyst wall include the following. While cyst walls of *Entamoeba* and *Giardia* have a single layer, a single glycopolymer (chitin and β-1,3-linked-GalNac, respectively), and small sets of one or two lectins, *A. castellanii* cyst walls have two layers and ostioles, at least two glycopolymers, and three large sets of lectins (31, 56, 57, 63, 64). Although cellulose and chitin synthases are each single copy, it will be difficult to knock out sets of genes encoding CWPs, even with the efficient gene editing methods available in other protists (27-29, 65, 66).

### Familiarity and novelty in *A. castellanii* CWPs

Although we expected Luke lectins with two or three CBM49s would be present in the cellulose-rich cyst wall, we could not have predicted the other abundant CWPs, because the 8-Cys domains of Leo lectins are unique to *Acanthamoebae* and the CAA domains of Jonah lectins were previously uncharacterized (27, 42, 49). While Luke lectins have two or three CBM49s, *D. discoideum* has dozens of proteins with a single CBM49 (Fig. S6) (44). The Luke lectin binds cellulose and chitin, while the *D. discoideum* proteins with a single CBM49 bind cellulose (46). Chitin-binding by DdCBM49 or SlCBM49 was not tested, because this glycopolymer is not present in *D. discoideum* and tomato walls. Demonstration that CBM49s of the Luke lectin also bind chitin fibrils is new, but is consistent with recent studies showing CBMs may bind more than one glycopolymer (67). The metalloprotease fused to an N-terminal CBM49 of *A. castellanii* is absent in *D. discoideum*, while *D. discoideum* adds two CBM49s to a cysteine proteinase, which lacks these domains in *A. castellanii*. The CBM49 may act to localize the metalloproteases to the *A. castellanii* cyst wall, as is the case for the chitin-binding domain in *Entamoeba* Jessie lectins or glucan-binding domain in *Toxoplasma* glucanases (56, 58). Alternatively, the CBM49 may suggest the metalloprotease cleaves glycopeptides rather than peptides. While the GH5 glycoside hydrolases of *A. castellanii* lack CBM49s, CBM49 is present at the C-terminus of GH9 glycoside hydrolases of *D. discoideum* and *S. lycopersicum* (27, 42, 44).

Even though *A. castellanii* Leo lectins and *E. histolytica* Jacob lectins share no common ancestry, they have 8-Cys and 6-Cys lectin domains, respectively, often separated by low complexity sequences (Fig. S7) (63, 64). *E. histolytica* low complexity sequences vary from strain to strain, contain cryptic sites for cysteine proteases, and are extensively decorated with *O*-phosphate-linked glycans (68). We have not yet identified any Asn-linked or *O*-linked glycans on Leo lectins or any of the other CWPs, but we expect they will be there. *A. castellanii* and oomycetes (*Pyromyces* and *Neocallmistic*) each contain proteins with arrays of CAA domains, but the sequences of the CAAs are so different that it is likely that concatenation of domains occurred independently (Fig. S8) (43, 44). Although *A. castellanii* is exposed to collagen in the extracellular matrix of the cornea, the protist lacks a homolog of the collagen-binding domain that is adjacent to the CAA domain in the *Bacillus anthracis* collagen-binding protein (49).

Concatenation of carbohydrate-binding domains in Luke, Leo, and Jonah lectins, which has previously been shown in WGA, Jacob lectins of *E. histolytica*, and peritrophins of insects, most likely increases the avidity of the lectins for glycopolymers (31, 63, 64, 69). The large number of genes encoding Luke, Leo, and Jonah lectins may be necessary to increase the quantity of CWPs coating glycopolymers in the cyst wall. For example, Luke(2) and Luke(3) lectins with two or three CBM49s, respectively, localize to the same place in mature cyst walls. Alternatively, there may be differences in timing and localization of CWPs within the same family, based upon a TKH-rich spacer in Leo lectins or transmembrane helices in Jonah lectins with three CAA domains (Fig. S2). Finally, other candidate CWPs, which are abundant but present at lower copy numbers (e.g. laccase or ferritin-domain protein) and so not tested here, may have important functions in the cyst wall.

### Possible translation of these results to the clinic

While the focus here was on the biochemistry and cell biology of the cyst wall, Luke, Leo, and Jonah lectins, which are conserved in all strains of *A. castellanii*, appear to be excellent targets for diagnostic antibodies, as has been shown for *Giardia* CWP1 and *Entamoeba* Jacob2 lectin (70, 71). Evidence for chitin in the cyst wall, admittedly not extensively developed here, suggests the possibility that chitin synthase inhibitors or chitinases might be used as therapeutics (72). Other investigators have explored the possibility of cellulose synthase inhibitors or cellulases as therapeutics versus *A. castellanii* cysts (73-76). The SIM methods, which worked so well here to study *A. castellanii* cyst walls, might be used to study the fine structure and assembly of other protist walls. Insights from studies of the *A. castellanii* cyst wall, which is relatively simple, may inform studies of fungal and plant walls, which are highly complex.

## MATERIALS AND METHODS

### Culture of trophozoites and preparation of encysting organisms and cysts

Culture and manipulation of *A. castellanii* were approved by the Boston University Institutional Biosafety Committee. *A. castellanii* Neff strain trophozoites were purchased from the American Type Culture collection. Trophozoites of *A. castellanii* MEEI 0184 strain, which was derived from a human corneal infection, were obtained from Dr. Noorjahan Panjwani of Tufts University School of Medicine (9). Trophozoites were grown in T-75 tissue culture flasks at 30°C in 10 ml ATCC medium 712 (PYG plus additives) (Sigma-Aldrich Corporation, St. Louis, MO) (11). Flasks containing log-phase trophozoites (free of cysts that form spontaneously in stationary cultures) were either chilled or scraped with a cell scraper to release adherent amoebae, which were concentrated by centrifugation at 500 × g for 5 min and washed twice with phosphate buffered saline (PBS). Approximately 10^7^ amoebae obtained from a confluent flask were induced to encyst by incubation at 30°C on non-nutrient agar plates (10). After 3 to 6 days incubation, 15 ml of PBS was added to agar plates, which were incubated on a shaker for 30 min at room temperature (RT). Encysting organisms were removed using a cell scraper and concentrated by centrifugation at 1,500 × g for 10 min.

### Preparation of mature cyst walls for SIM, TEM, and mass spectrometry

Mature cysts were washed in PBS and suspended in lysis buffer (10 mM HEPES, 25 mM KCl, 1 mM CaCl_2_, 10 mM MgCl_2_, 2% CHAPS, and 1X Roche protease inhibitor) (Sigma-Aldrich). For SIM, cysts in 500-µl lysis buffer were broken four times for 2 min each with 200 µl of 0.5 mm glass beads in a Mini-Beadbeater-16 (BioSpec Products, Bartlesville, OK). For TEM, where glass beads cannot be used, cysts in 200-µl lysis buffer were broken by sonication four times for 20 seconds each in continuous mode in a Sonicator Cell Disruptor (formerly Heat Systems Ultrasonic, now Qsonica, Newtown, CT). Broken cysts were added to the top a 15-ml falcon tube containing 60% sucrose and centrifuged at 4,000 × g for 10 min. The broken cyst wall pellet was suspended in PBS buffer and washed three times at 10,000 × g in a microcentrifuge. The cyst wall pellet was used directly for labeling for SIM or fixation for TEM. For mass spectrometry, the cyst wall pellet broken in the bead beater was overlaid on gradient containing 2 ml each of 20%, 40%, 60% and 80% Percoll (top to bottom) and centrifuged for 20 min at 3,000 × g. The layer between 60% and 80% Percoll, where the broken cyst walls were located, was collected and washed in PBS. The cyst wall preparation was suspended in 10 ml of PBS, placed in a syringe, and forced through a 25-mm diameter Whatman Nuclepore Track-Etched Membrane with 8-µm holes (Sigma-Aldrich). The cellular debris, which passed through the membranes, was discarded. The membrane was removed from the cassette, suspended in 5 ml of PBS, and vortexed to release cyst walls. The membrane was removed, and cyst walls were distributed in microfuge tubes and pelleted at 15,000 × g for 10 min. The pellet was suspended in 50 µl PBS and stored at −20°C prior to trypsin digestion and mass spectrometry analysis.

### SIM of glycopolymers of mature cysts and purified cyst walls

A GST-AcCBM49 fusion-construct, which contains the N-terminal CBM49 of a representative Luke lectin minus the signal peptide, was prepared by codon optimization (76 to 330-bp coding region of ACA1_377670) (GenScript, Piscataway, NJ). It was cloned into pGEX-6p-1 (GE Healthcare Life Sciences, Marlborough, MA) for cytoplasmic expression in BL21(DE3) chemically competent *E. coli* (Thermo Fisher Scientific, Waltham, MA) (41, 42). Expression of GST-AcCBM49 and GST were induced with 1 mM IPTG for 4 hr at RT, and GST-fusions were purified on glutathione-agarose and conjugated to Alexa Fluor 594 succinimidyl esters (red) (Molecular Probes, Thermo Fisher Scientific). Approximately 10^6^ mature cysts or cyst walls were washed in PBS and fixed in 1% paraformaldehyde buffered with phosphate for 15 min at RT. Pellets were washed two times with Hank’s Buffered Saline Solution (HBSS) and incubated with HBSS containing 1% bovine serum albumin (BSA) for 1 hour at RT. Preparations were then incubated for 2 hr at 4°C with 2.5 µg GST-CBM49 conjugated to Alexa Fluor 594 and 12.5 µg of WGA (Vector Laboratories, Burlingame, CA) conjugated to Alexa Fluor 488 in 100 µl HBSS (37, 38). Finally, pellets were labeled with 100 µg of calcofluor white M2R (Sigma-Aldrich) in 100 µl HBSS for 15 min at RT and washed five times with HBSS (39, 40). Preparations were mounted in Mowiol mounting medium (Sigma-Aldrich) and observed with widefield and differential interference contrast microscopy, using a 100x objective of a Zeiss AXIO inverted microscope with a Colibri LED (Carl Zeiss Microcopy LLC, Thornwood, NY). Images were collected at 0.2-μm optical sections with a Hamamatsu Orca-R2 camera and deconvolved using ZEN software (Zeiss). Alternatively, SIM was performed with a 63-x objective of a Zeiss ELYRA S.1 microscope at Boston College (Chestnut Hill, MA), and 0.09-μm optical sections deconvolved using Zen software (30).

### TEM of intact and purified walls

High-pressure freezing and freeze substitution were used to prepare cyst and cyst walls for TEM at the Harvard Medical School Electron Microscope facility (35). To make them noninfectious, we fixed mature cysts in 1% paraformaldehyde for 10 min at RT and washed them 2 times in PBS. Cyst walls in PBS were pelleted, placed in 6-mm Cu/Au carriers, and frozen in an EM ICE high-pressure freezer (Leica Microsystems, Buffalo Grove, Il). Freeze substitution was performed in a Leica EM AFS2 instrument in dry acetone containing 1% _dd_H_2_0, 1% OsO_4_, and 1% glutaraldehyde at −90°C for 48 hr. The temperature was increased 5°C/hour to 20°C, and samples were washed 3 times in pure acetone and once in propylene oxide for 10 min each. Samples were infiltrated with 1:1 Epon:propylene oxide overnight at 4°C and embedded in TAAB Epon (Marivac Canada Inc. St. Laurent, Canada). Ultrathin sections (80 to 100 nm thick) were cut on a Leica Reichert Ultracut S microtome, picked up onto copper grids, stained with lead citrate, and examined in a JEOL 1200EX transmission electron microscope (JEOL USA, Peabody, MA). Images were recorded with an AMT 2k CCD camera.

### Mass spectrometry of tryptic and chymotryptic peptides from cyst walls

Broken cyst walls, prepared as above, were dissolved into 50 mM NH_4_HCO_3_, pH 8.0, reduced with 50 mM dithiothreithol (DTT) for 20 min at 60°C, alkylated with iodoacetamide (IAA) for 20 min at RT, and then digested with proteomics grade trypsin (Sigma-Aldrich) overnight at 37°C. Alternatively broken cyst walls either before or after digestion with trypsin were reconstituted in 1× reducing SDS/PAGE loading buffer and run on a 4–20% precast polyacrylamide TGX gel (Bio-Rad). Bands stained by colloidal Coomassie blue were excised and washed with 50 mM NH_4_HCO_3_/acetonitrile (ACN). Reduction, alkylation, and trypsin/chymotrypsin digestion were performed in-gel. Peptides were dried and desalted using C18 ZipTip concentrators (EMD Millipore). Peptides from five biological replicates for both *in solution* and *in-situ* hydrolysis were dissolved in 2% ACN, 0.1% formic acid (FA) and separated using a nanoAcquity-UPLC system (Waters) equipped with a 5-μm Symmetry C18 trap column (180 μm × 20 mm) and a 1.7-μm BEH130 C18 analytical column (150 μm × 100 mm). Samples were loaded onto the precolumn and washed for 4 min at a flow rate of 4 μl/min with 100% mobile phase A (99% Water/1% ACN/0.1% FA). Samples were eluted to the analytical column with a gradient of 2– 40% mobile phase B (99% ACN/1% Water/0.1% FA) delivered over 40 or 90 min at a flow rate of 0.5 μl/min. The analytical column was connected online to a QE or a QE-HF Mass Spectrometer (Thermo Fisher Scientific) equipped with a Triversa NanoMate (Advion Inc., Ithaca, NY) electrospray ionization (ESI) source, which was operated at 1.7 kV. Data were acquired in automatic Data Dependent top 10 (QE) or top 20 (QE-HF) mode. Automated database searches were performed using the PEAKS software suite version 8.5 (Bioinformatics Solutions Inc., Waterloo, ON, Canada). The search criteria were set as follows: trypsin/chymotrypsin as the enzyme with ≤ 3 missed cleavages and ≤ 1 non-specific cleavage, the error tolerances for the precursor of 5 ppm and 0.05 Da for fragment ions, carbamidomethyl cysteine as a fixed modification, oxidation of methionine, Pyro-glu from glutamine, and deamidation of asparagine or glutamine as variable modifications. The peptide match threshold (−10 logP) was set to 15, and only proteins with a minimum of two unique peptides were considered. The mass spectrometry proteomics data have been deposited to the ProteomeXchange Consortium (http://proteomecentral.proteomexchange.org) via the PRIDE partner repository with the dataset identifiers PXD011826 and 10.6019/PXD011826 (77).

### Bioinformatic characterization of candidate CWPs

Signal peptides and transmembrane helices were predicted using SignalP 4.1 and TMHMM, respectively (45, 48). Glycosylphosphatidylinositol anchors were searched for using big-PI (47). AmoebaDB, which contains sequence information from the Neff strain and ten other *Acanthamoeba* strains, was used to identify genome sequences, predict introns, and identify paralogous proteins (28, 29). The NR database at the NCBI was used to identify homologs of candidate CWPs in other species and to identify conserved domains (43). Carbohydrate-binding modules were searched using CAZy and InterPro databases (27, 44).

### Expression and visualization of GFP-fusions in transfected *A. castellanii*

We used RT-PCR from RNA of encysting protists to obtain the coding sequences of a representative Luke(2) lectin (840-bp CDS of ACA1_377670), Luke(3) lectin (1293-bp CDS of ACA1_245650), Leo lectin (562-bp CDS of ACA1_074730), and Jonah(1) lectin (1596-bp CDS of ACA1_164810) (Figs. 3 and S2). Please see Supplemental Table 1 for a list of primers used to make all the constructs. Using NEBuilder HiFi DNA assembly (New England Biolabs, Ipswich, MA), we cloned each CDS into the pGAPDH plasmid, which was a kind gift from Yeonchul Hong of Kyongpook National University School of Medicine, Deagu, Korea (33). pGAPDH contains a neomycin resistance gene under a TATA-box promoter (for selection with G418) and a glyceraldehyde 3-phosphate dehydrogenase promoter for constitutive expression of GFP-fusions (Fig. S1 and Supplemental Table 1). The GFP tag was placed at the C-terminus of each CWP, and a polyadenylation sequence was added downstream of the GFP-fusion’s stop codon. For expression of CWP genes under their own promoters, we replaced the GAPDH promoter with 446-bp from the 5 ‘UTR of the Luke(2) gene, 500-bp from the 5’ UTR of the Luke(3) gene, 486-bp from the 5’ UTR of the Leo gene, and 571-bp of the 5’UTR of the Jonah(1) gene, each cloned from the genomic DNA.

As a control, SP-GFP, which contains a 60-bp sequence encoding an N-terminal signal peptide of Luke(2) lectin, was expressed behind a GAPDH promoter. As another control, the 470-bp 5’ UTR and 525-bp CDS of CSP21 (ACA1_075240) was made with a GFP tag (22).

Transfections in *A. castellanii* were performed as described previously (32, 33) with some modifications. Briefly, 5 × 10^5^ log-phase trophozoites were allowed to adhere to 6-well plates in ATCC medium 712 for 30 min at 30°C. The adherent trophozoites were washed and replaced with 500 µl of non-nutrient medium (20 mM Tris-HCl (pH 8.8), 100 mM KCl, 8 mM MgSO_4_, 0.4 mM CaCl_2_ and 1 mM NaHCO_3_). In an Eppendorf tube, 4 µg of Midiprep (PureLink HiPure Midiprep Kit, Thermo Fisher Scientific) plasmid DNA was diluted to 100 µl with non-nutrient medium. Twenty microliters of SuperFect Transfection Reagent (Qiagen Inc, Germantown, MD) was added to the DNA suspension, mixed gently by pipetting five times, and incubated for 10 min at RT. Six hundred microliters of non-nutrient medium were added to the DNA-SuperFect mix, and the entire suspension was added to the trophozoites adhering to the 6-well culture plate. The culture plate was incubated for 3 hr at 30°C, after which the non-nutrient medium was replaced with ATCC medium 712 and incubated for another 24 hr at 30°C. To select for transfectants, we added 12.5 µg/ml of Gibco G418 antibiotic (Thermo Fisher Scientific) to the culture after 24 hr, and we changed the medium plus antibiotic every 4 days. After 2 to 4 weeks, the transfectants were growing robustly in the presence of the antibiotic, and trophozoites and/or cysts expressing GFP were detected by widefield microscopy. These organisms were induced to encyst, fixed, labeled with WGA and CFW, and examined by widefield microscopy and SIM, as described above.

### Binding of MBP-CWP fusions to microcrystalline cellulose and chitin beads

MBP-fusion constructs were prepared by cloning the cDNA of a representative Luke(2) lectin (60 to 843-bp CDS of ACA1_377670) and a representative Jonah(1) lectin (70 to 1599-bp CDS of ACA1_164810) without their signal sequences into pMAL-p2x vector (New England Biolabs) for periplasmic expression in BL21-CodonPlus(DE3)-RIPL (Agilent Technologies, Lexington, MA) (34). For the MBP-fusion, the Leo CDS without the signal sequence (67 to 564-bp of ACA1_074730) was codon optimized (GenScript) and cloned into pMAL-p2x vector. MBP-Luke(2) was induced with 250 µM IPTG for 5 hr at RT; MBP-Jonah(1) was induced with 1 mM IPTG for 5 hr at RT; and MBP-Leo was induced with 250 µM IPTG for 3.5 hr at 37°C. MBP-fusion proteins were purified with amylose resin following the manufacturer’s instructions (GE Healthcare, Pierce, Agilent Technologies and New England Biolabs). MBP-fusions (1 µg each in 100 µl of 1% NP40) were incubated with 0.5 µg Avicel microcrystalline cellulose (Sigma-Aldrich) or a 50-µl slurry of magnetic chitin beads (New England Biolabs) for 3 hr at 4°C with rocking. Cellulose was centrifuged, and magnetic chitin beads were placed in a magnet to collect the supernatant (unbound fraction) and pellet (bound fraction). The pellet was washed three times with 1% NP40. To solubilize proteins, the input material (total), unbound, and bound fractions were boiled in SDS sample buffer. MBP-proteins were separated on SDS-PAGE gels, blotted to PVDF membranes, blocked in 5% BSA, and detected using anti-MBP antibodies (New England Biolabs).

### Western blots of *A. castellanii* trophozoite and cyst lysates probed with anti-lectin rabbit antibodies

Log-phase trophozoites and 36-hr-old cysts were harvested, and the total protein solubilized in SDS sample buffer, run in SDS-PAGE gels, blotted on PVDF membranes, and blocked in 5% BSA. MBP-CWP fusions and MBP alone were run in adjacent lanes as positive and negative controls, respectively. The blots were probed with 1:100 dilutions of rabbit polyclonal antibodies (Li International) raised to 16- or 50-amino acid peptides of representative Luke(2) lectin (residues 230-279 of ACA1_377670), Leo lectin (residues 124-139 of ACA1_074730) and Jonah(1) lectin (residues 362-411 of ACA1_164810) (Fig. 3). A 1:1000 dilution of anti-rabbit IgG-HRP (BioRad) was used as secondary antibody and Super Signal West Pico PLUS (ThermoFisher Scientific) for chemiluminescent detection. Coomassie stained gels were run in parallel for loading control.

## ACKNOWLEDGMENTS

We thank Maria Ericsson of Harvard University for help with transmission electron microscopy. We thank Bret Judson and Marc-Jan Gubbels of Boston College for help with structured illumination microscopy, which was supported by NSF grant No. 1626072. This work was also supported by awards from the National Institute of Allergy and Infectious Diseases of the NIH to J.S. (R01 AI110638), from the National Institute of General Medical Science of the NIH to C.E.C. (P41 GM104603), and from the Department of Energy to B.R.U. (DESC0015662). We also thank Dean Jeffrey Hutter of the Boston University Goldman School of Dental Medicine for his support of research in the Samuelson lab.

## Supplemental files include eight figures, two Excel files, and one table

**FIG S1.**
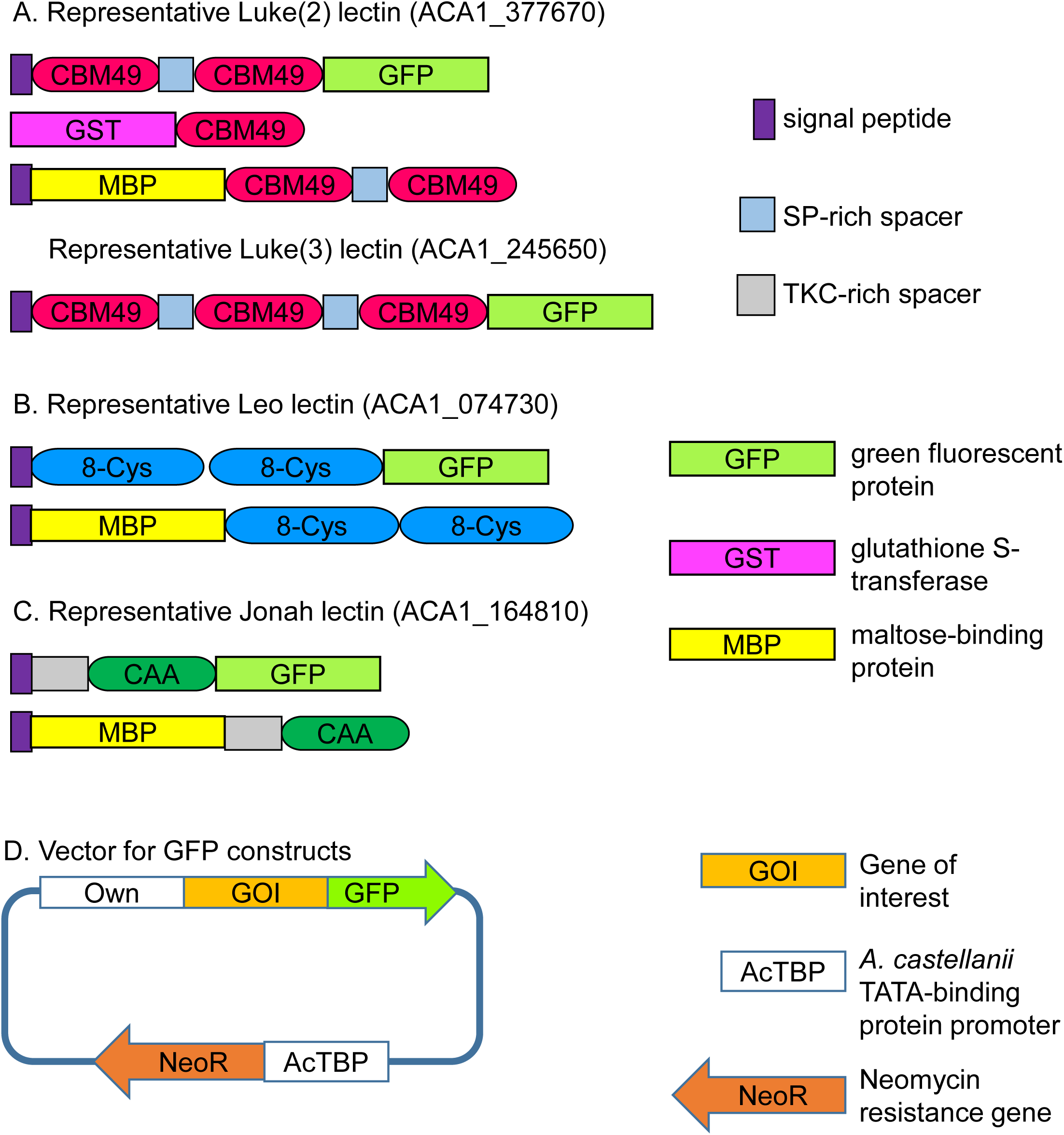
Constructs made to localize CWPs Ac cyst wall and to determine their binding to microcrystalline cellulose and chitin beads. A. A representative Luke(2) lectin with two CBM49s was used to make GFP-, GST-, and MBP-fusions. A representative Luke(3) lectin with three CBM49s was used to make a GFP-fusion protein. B. A representative Leo lectin was made into GFP- and MBP-fusions. C. A representative Jonah(1) lectin with a single CAA domain was made into GFP- and MBP-fusions. D. Vectors for expressing GFP-fusions in transfected *A. castellanii* under its own promoter contain a neomycin resistance gene under a TATA-binding protein promoter (32, 33). Primers for making constructs are listed in Supplemental Table 1.

**FIG S2.**
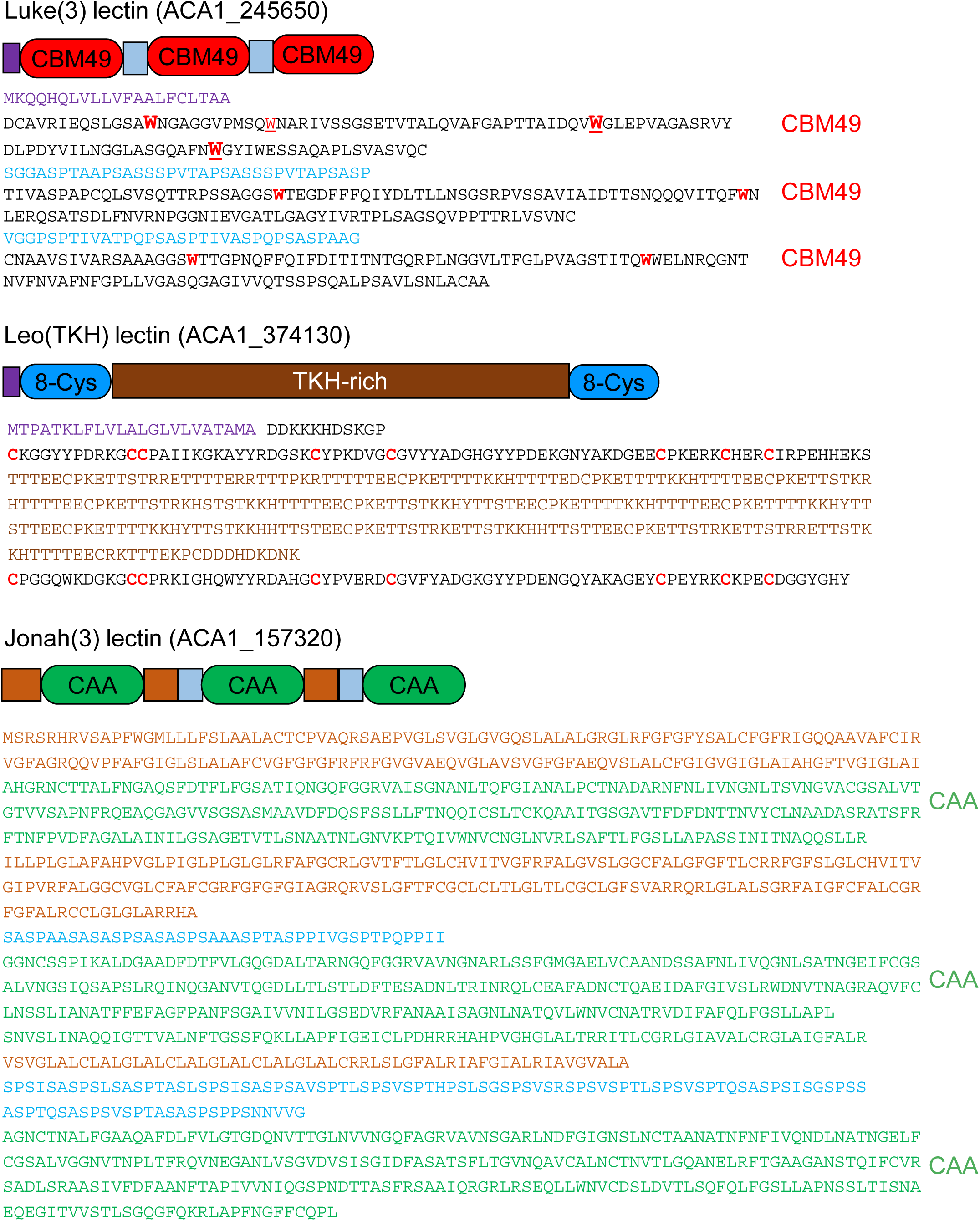
Sequences of candidate CWPs, which differ in at least one essential property from Luke, Leo, and Jonah lectins used for localization and binding studies. A Luke(3) lectin has an N-terminal signal peptide (purple) and three CBM49s separated by short Ser- and Pro-rich spacers (light blue). The CBM49s contain conserved Trp (red Ws) present in the representative Luke lectin (Fig. 3). A Leo(TKH) lectin has a signal peptide, two domains containing eight Cys residues each (red Cs), and a long Thr-, Lys-, and His-rich spacer (brown). A Jonah(3) lectin has three CAA domains (green), hydrophobic regions (tan), and short Ser- and Pro-rich spacers (light blue).

**FIG S3.**
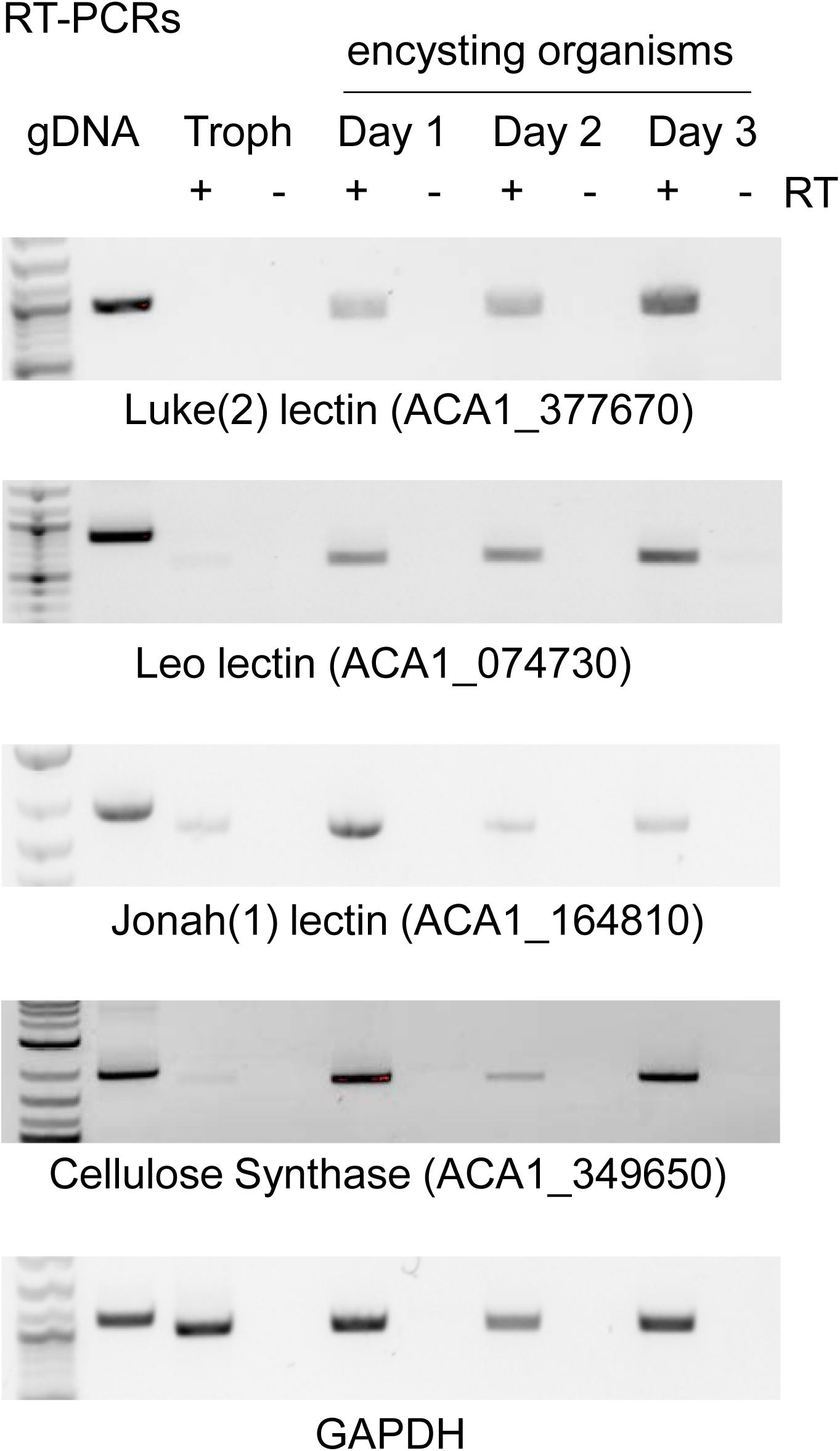
RT-PCR shows mRNAs of representative Luke, Leo, and Jonah lectins, as well as those of cellulose synthase, are encystation-specific. DNA and total RNA were extracted from trophozoites and organisms encysting for one to three days. RT-PCRs were performed with primers specific for segments of each CWP mRNA, as well as primers specific for segments of mRNAs for GAPDH and cellulose synthase (see Supplemental Table 1). PCR with DNA was used as a positive control, while omission of reverse-transcriptase (-RT) was used as a negative control. Messenger RNAs encoding CWPs and cellulose synthase were absent or nearly absent in trophozoites but were easily detectable in encysting organisms. In contrast, mRNAs for GAPDH were expressed by both trophozoites and encysting organisms (33).

**FIG S4.**
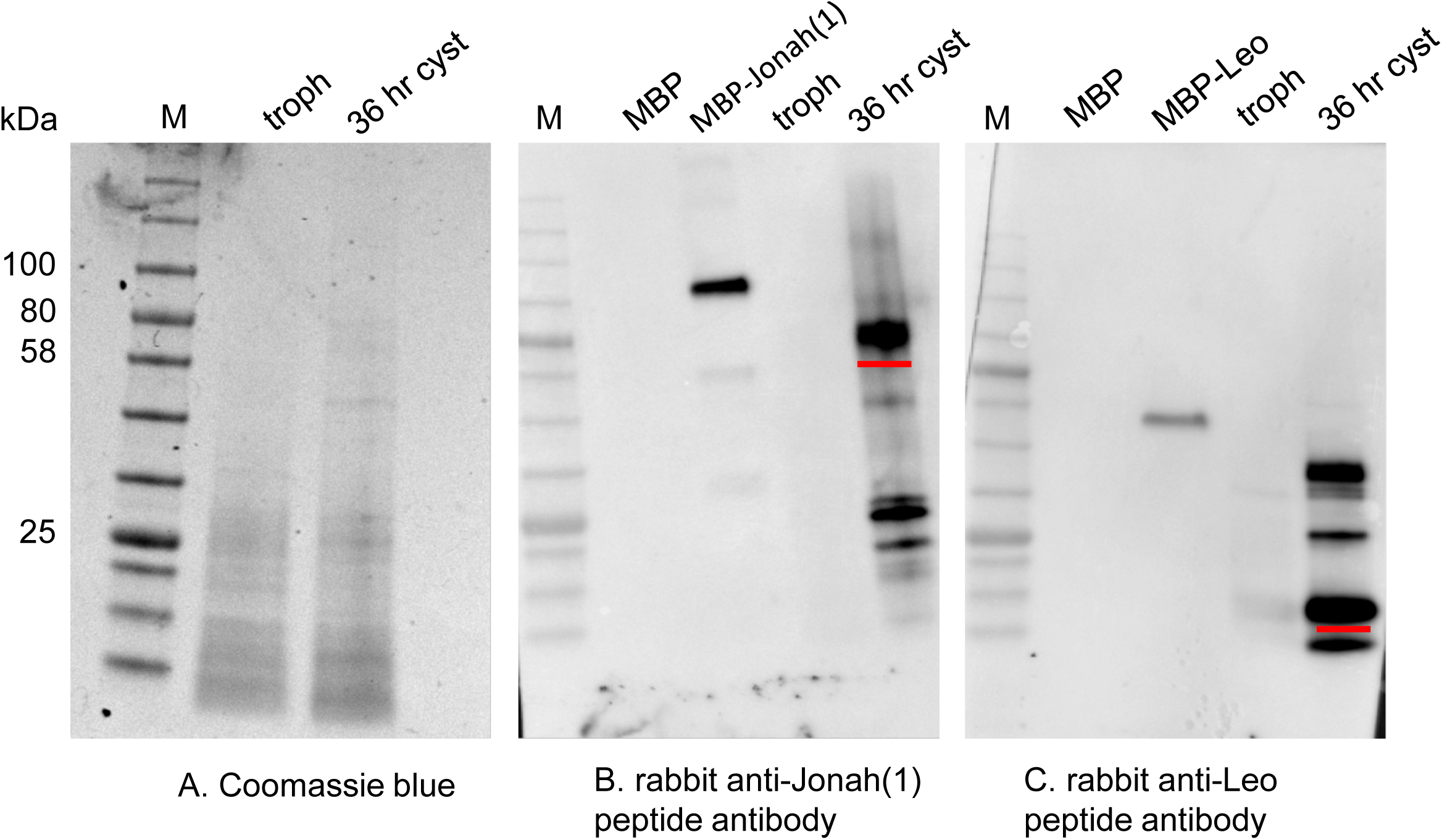
Western blots with rabbit antibodies to peptides of Jonah and Leo lectins show each CWP is absent in trophozoites but is easily detected in mature cysts. A. Coomassie blue stain of proteins of lysed trophozoites and cysts, as well as molecular weight standards (M). B. Western blotting showed rabbit antibodies to a 50-amino acid peptide of a representative Jonah(1) lectin (underlined in Fig. 3) bound to a cyst protein of the predicted size (red underline) and to an MBP-Jonah(1) fusion-protein made in the periplasm of bacteria. The antibody also bound to degradation products of Jonah(1) lectin. In contrast, the anti-Jonah(1) antibody did not bind to either trophozoites or MBP alone (negative controls). C. Rabbit antibodies to a 16-amino acid peptide of a representative Leo lectin also bound to cyst proteins and to an MBP-Leo fusion but not to trophozoite proteins or to MBP alone. In addition to Leo of the predicted size (red underline), anti-Leo antibodies to a higher molecular weight form, which may be a dimer. These results confirmed encystation-specific expression of Jonah(1) and Leo lectins (Figs. 4 and S3). None of the rabbit antibodies the CWP peptides were useful for labeling cyst walls for widefield microscopy or SIM.

**FIG S5.**
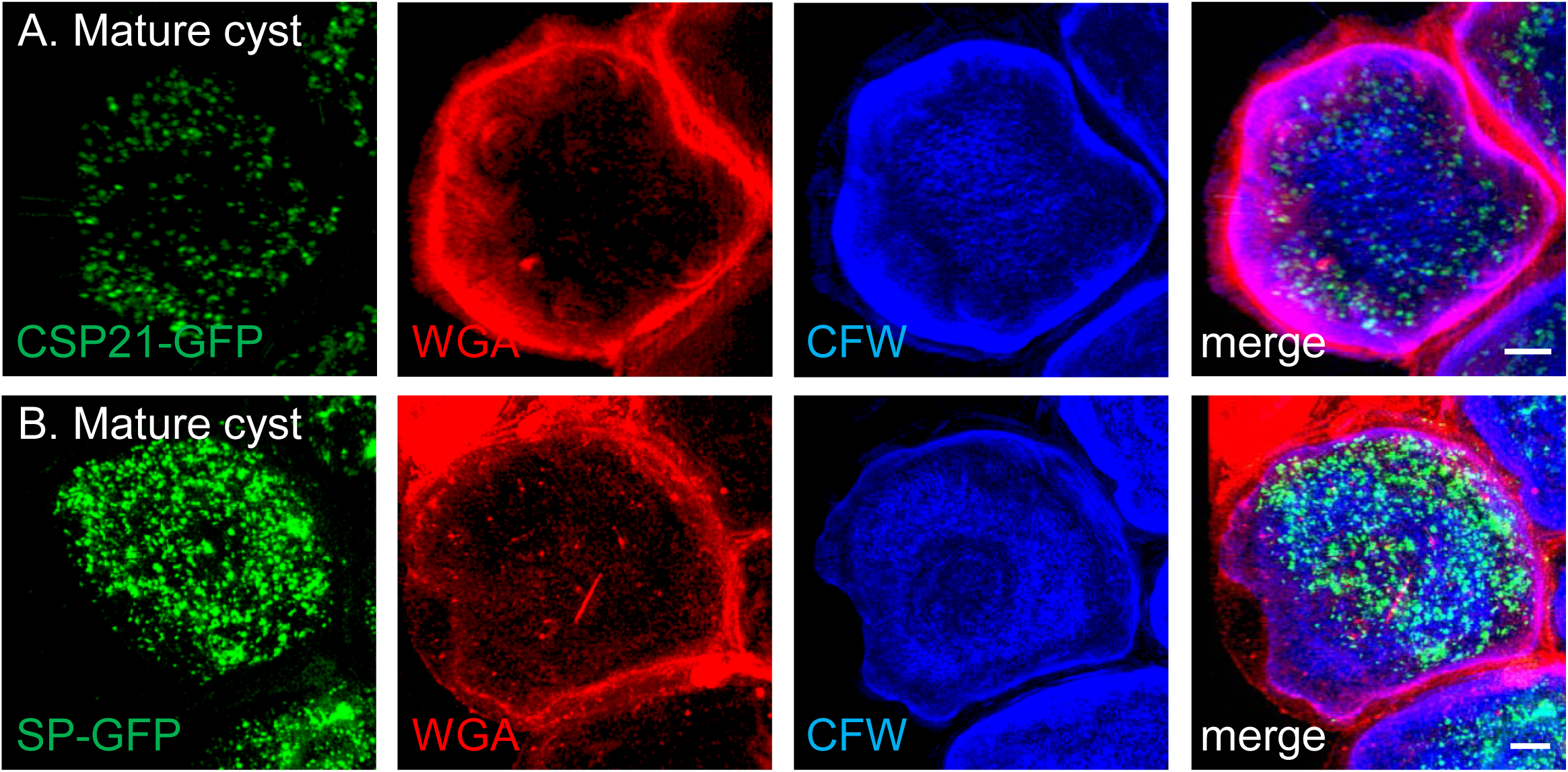
Control GFP constructs localize to the cytosol (CSP21-GFP) and secretory vesicles (GFP with an N-terminal signal peptide, SP-GFP) of mature cysts. A. The 21-kDa cyst-specific protein (CSP21) fused to GFP was absent in trophozoites (not shown) but formed punctate structures in the cytosol of cysts (22). B. GFP with an N-terminal signal peptide from Luke lectin and expressed under a GAPDH promoter localized to secretory vesicles of mature cysts (33). These controls make it unlikely that localizations of CWP-GFP constructs to the cyst wall were artifacts (Fig. 4).

**FIG S6.**
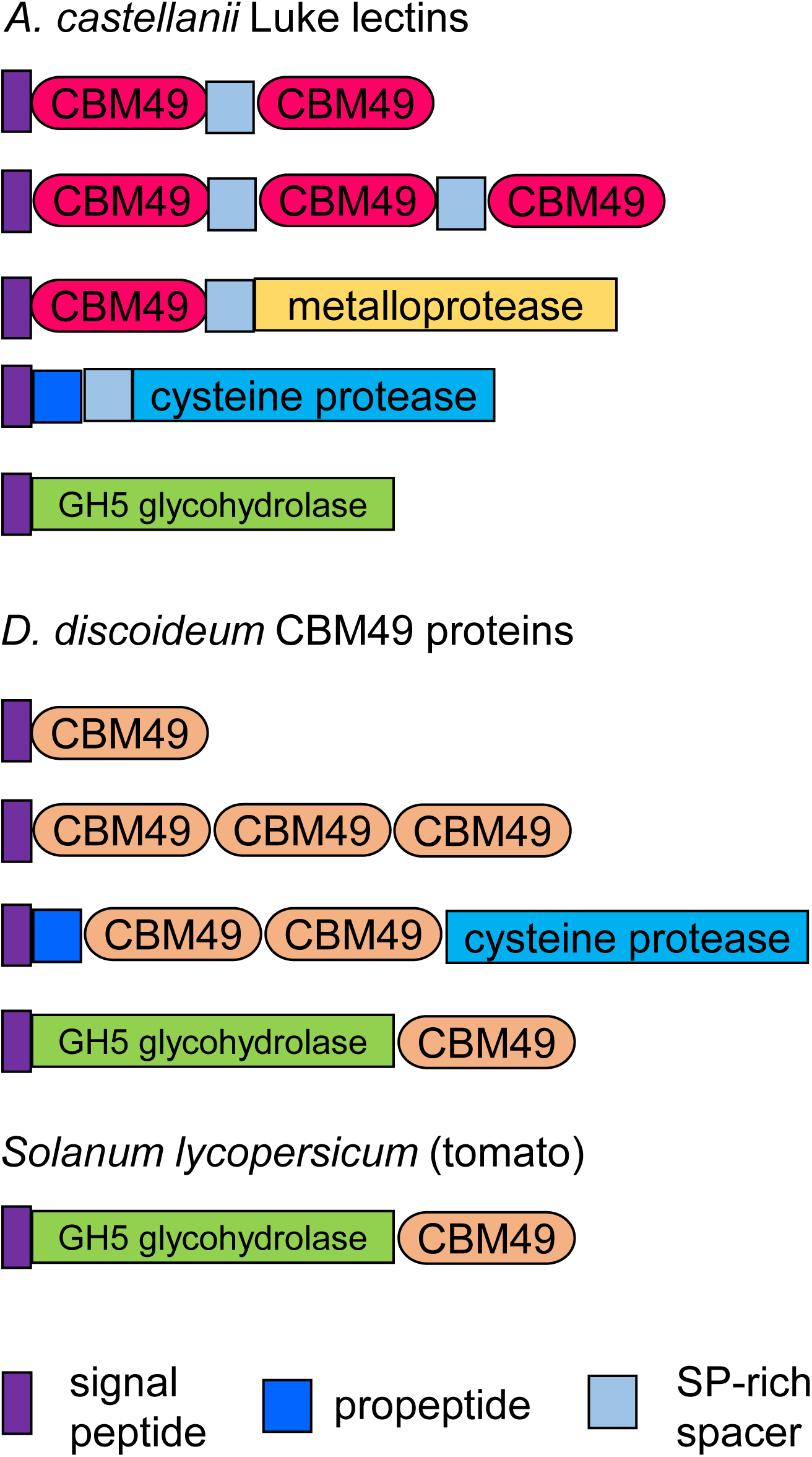
Contrasting uses of CBM49 by *A. castellanii* and *D. discoideum*. CBM49, which was first shown to be a cellulose-binding domain at the C-terminus of tomato cellulase, is repeated two or three times in Luke lectins of *A. castellanii* and is also present at the N-terminus of a metalloprotease. In contrast, CBM49 is present in a single copy in the majority of *D. discoideum* proteins and as three copies in rare proteins. CBM49 is also present in two copies in a *D. discoideum* cysteine protease and as a single copy in a GH5 glycoside hydrolase.

**FIG S7.**
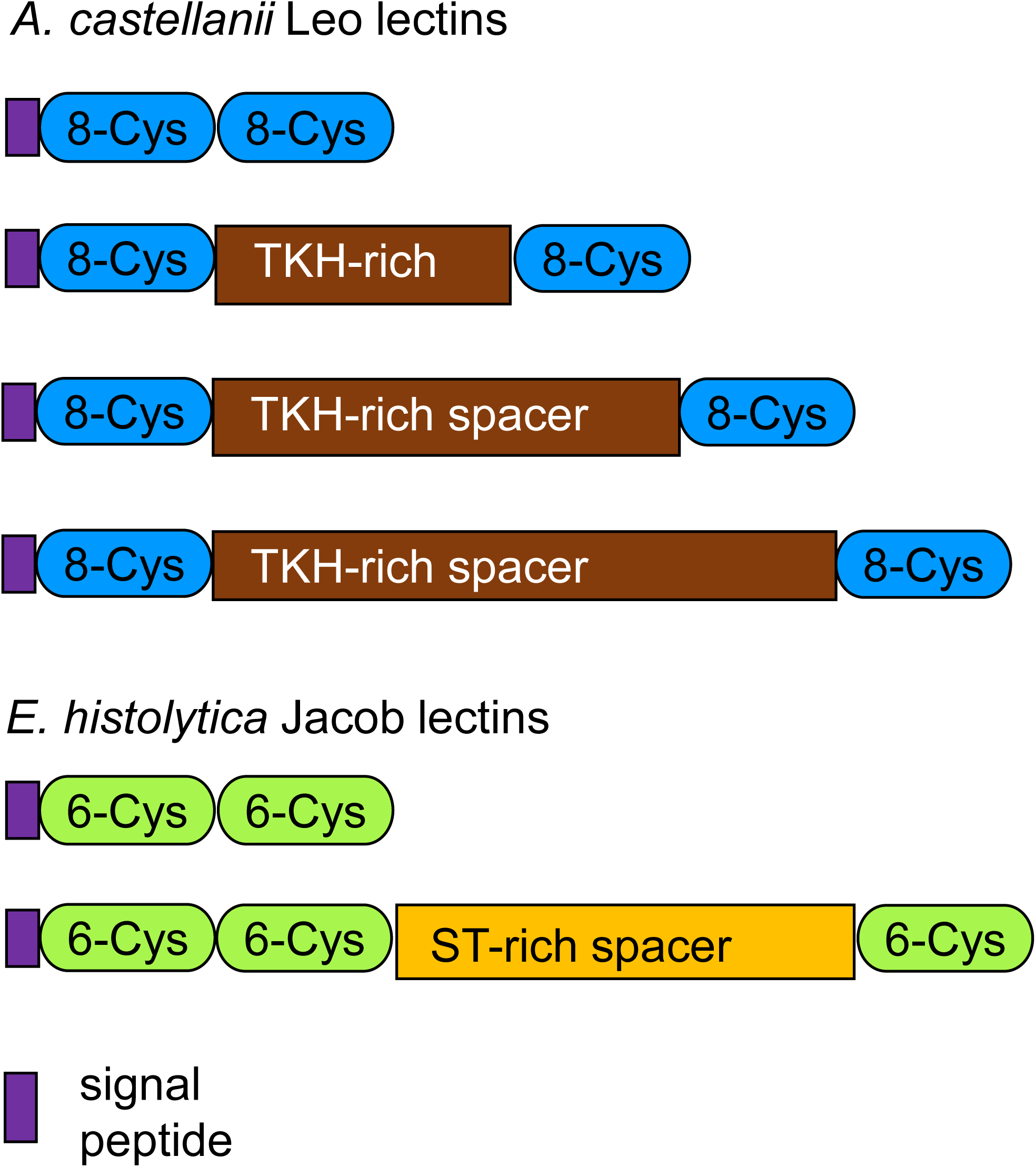
*A. castellanii* Leo lectins and *E. histolytica* Jacob lectins have common structures, even though they share no common ancestry (convergent evolution). Abundant cyst wall proteins of *A. castellanii* (Leo lectins) and *E. histolytica* (Jacob lectins) have unique 8-Cys or 6-Cys domains, respectively, that bind cellulose or chitin. In each protist, some of the lectins lack spacers, while others have spacers rich in Thr, Lys, and His (*A. castellanii*) or Ser and Thr (*E. histolytica*).

**FIG S8.**
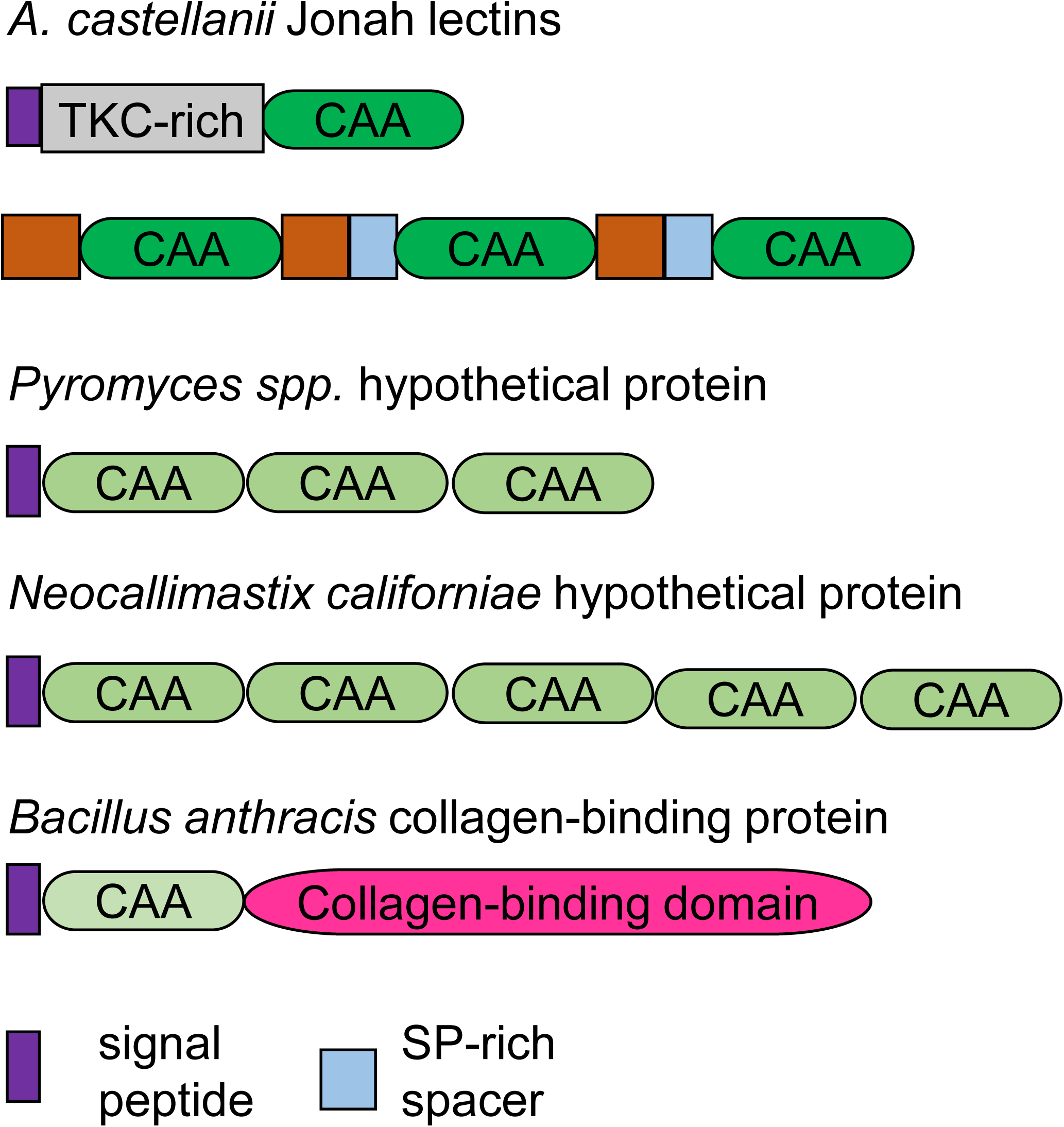
Contrasting use of choice of anchor A (CAA) domains in *A. castellanii*, oomycetes, and bacteria. Jonah lectins, which are abundant in cyst walls of *A. castellanii*, have one or three CAA domains. The former are preceded by Thr-, Lys-, and Cys-rich sequences (gray), while the latter are separated by Ser- and Pro-rich spacers (blue) and hydrophobic domains (tan). Predicted proteins of oomycetes (*Pyromyces* or *Neocallimastix*) have three to five CAA domains, while the spore coat protein of *Bacilllus* has a single CAA domain attached to a collagen-binding domain, which is absent in *A. castellanii*.

**Excel file S1** This file lists the most abundant proteins identified by mass spectrometry of cyst walls purified on the Percoll gradient and retained on a membrane containing 8-µm pores. Proteins with <7 unique peptides have been removed, because they are dominated by cytosolic contaminants. Luke, Leo, and Jonah lectins have been highlighted in orange. Other candidate CWPs are marked in yellow.

**Excel file S2** Complete list of proteins identified by mass spectrometry of cyst walls. This list includes a preparation that was heavily contaminated with cytosolic proteins, because the Percoll gradient and porous membrane were omitted during their purification. Only proteins with at least two unique peptides are included.

**Supplemental Table 1.**
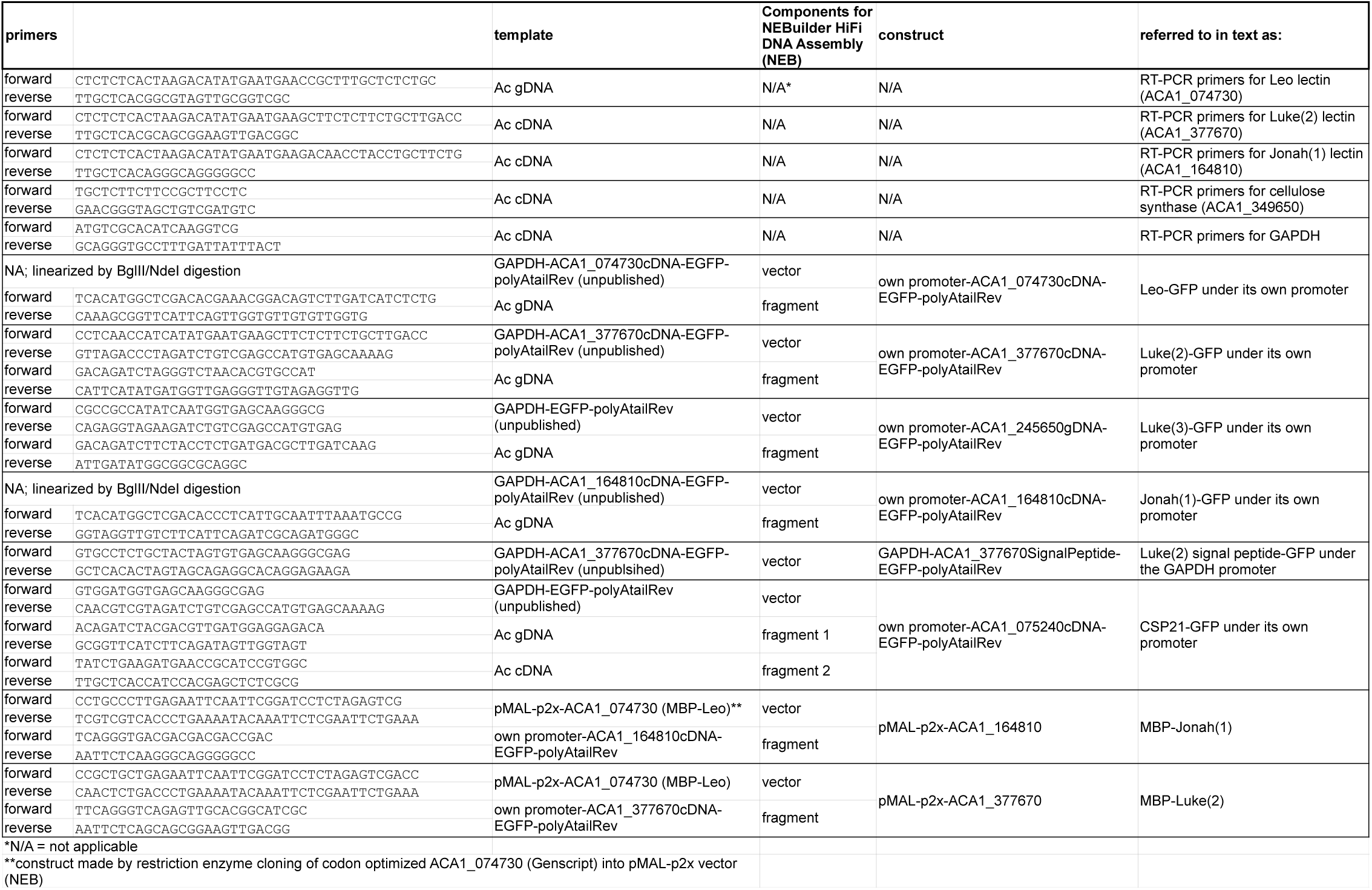
Primers used for RT-PCR and construction of GFP- and MBP-fusions.

